# A Safe Harbor-Targeted CRISPR/Cas9 Homology Independent Targeted Integration (HITI) System for Multi-Modality Reporter Gene-Based Cell Tracking

**DOI:** 10.1101/2020.02.10.942672

**Authors:** John J Kelly, Moe Saee-Marand, Nivin N Nyström, Yuanxin Chen, Melissa M Evans, Amanda M Hamilton, John A Ronald

**Author notes:** Address correspondence to: Dr. John A. Ronald, Robarts Research Institute, Schulich School of Medicine and Dentistry, University of Western Ontario, London, Ontario, N6A 5B7, Canada.

## Abstract

Imaging reporter genes can provide valuable, longitudinal information on the biodistribution, growth and survival of engineered cells in preclinical models and patients. A translational bottleneck to using reporter genes in patients is the necessity to engineer cells with randomly-integrating vectors. CRISPR/Cas9 targeted knock-in of reporter genes at a genomic safe harbor locus such as adeno-associated virus integration site 1 (AAVS1) may overcome these safety concerns. Here, we built Homology Independent Targeted Integration (HITI) CRISPR/Cas9 minicircle donors for precise AAVS1-targeted simultaneous knock-in of fluorescence, bioluminescence, and MRI (*Oatp1a1*) reporter genes. Our results showed greater knock-in efficiency at the AAVS1 site using HITI vectors compared to homology-directed repair donor vectors. Characterization of select HITI clones demonstrated functional fluorescence and bioluminescence reporter activity as well as significantly increased Oatp1a1-mediated uptake of the clinically-approved MRI agent gadolinium ethoxybenzyl diethylenetriamine pentaacetic acid. As few as 10^6^ Oatp1a1-expressing cells in a 50 µl subcutaneous injection could be detected *in vivo* with contrast-enhanced MRI. Contrast-enhanced MRI also improved the conspicuity of both sub-cutaneous and metastatic Oatp1a1-expressing tumours prior to them being palpable or even readily visible on pre-contrast images. Our work demonstrates the first CRISPR/Cas9 HITI system for knock-in of large DNA donor constructs at a safe harbor locus, enabling multi-modal longitudinal *in vivo* imaging of cells. This work lays the foundation for safer, non-viral reporter gene tracking of multiple cell types.

## Introduction

Molecular-genetic imaging with reporter genes permits the *in vivo* visualization and tracking of engineered cells, and thus, allows one to track the biodistribution, persistence, viability and in some cases, activation state, of such cells^1,2^. Several reporter genes currently exist for visualizing engineered cells using pre-clinical optical fluorescence imaging (FLI) and bioluminescence imaging (BLI)^3–5^ as well as those for clinical modalities such as magnetic resonance imaging (MRI), positron emission tomography (PET) and photoacoustic imaging^6–8^. These non-invasive cell tracking tools are invaluable for understanding mechanisms of disease progression and the evaluation of treatments in pre-clinical animal models. Important examples in cancer research include the tracking of therapeutic stem cells^9–11^, tracking immune cell migration, cancer progression and metastasis^12–14^, and evaluating tumour response to novel anticancer therapeutics^15,16^. More recently, the use of reporter genes to track therapeutic cells has been translated into the clinic. In this case, cytotoxic T cells were engineered to express a chimeric antigen receptor to target glioma cells, as well as a herpes simplex virus type 1 thymidine kinase (HSV1-TK) dual reporter-suicide gene (that selectively uptakes the PET tracer [^18^F]FHBG) to track the localization and viability of the injected therapeutic cells in glioma patients^17,18^.

Although reporter genes have great potential for therapeutic cell tracking, their functionality is best utilized when the genes are stably integrated into the desired cell’s genome – allowing reporter gene expression throughout the lifetime of the cell and in any subsequent daughter cells. Retroviral vectors, such as those derived from HIV lentiviruses, have generally been used for transgene integration due to their high transfection efficiency, large transgene capacity and their ability to transduce a variety of dividing and non-dividing cell types. However, the slow acceptance of using reporter genes for tracking cell-based therapies may in part be due to the increased risk of random or quasi-random insertional mutagenesis when transgenes are delivered using viral vectors^19^. Indeed, in previous clinical trials involving children with X-linked severe combined immunodeficiency, a Moloney murine leukemia virus–based γ-retrovirus vector expressing the interleukin-2 receptor γ-chain (γc) complementary DNA successfully restored immunity in most patients. However, 5 of the 20 patients also developed leukemia, of which one child died, as a result of insertional mutagenesis and transactivation of proto-oncogenes^20–22^. For future cell-based therapies, it is therefore highly desirable to edit cells with reporter genes in a safe and site-specific manner. The application of such editing tools would allow for longitudinal cell tracking to confirm that the cells are performing their intended role, as well as to detect any ectopic growths or misplaced targeting at the earliest time point. This will ultimately give the clinician greater control and confidence in the outcomes of the targeted therapy.

Genomic safe harbors can incorporate exogenous pieces of DNA and permit their predictable function, but do not cause alterations to the host genome, nor pose a risk to the host cell or organism^23^. Several studies have successfully used genome editing tools such as zinc finger nucleases (ZFN) and transcription activator-like effector nucleases (TALENs) to incorporate reporter genes at the adeno-associated virus integration site 1 (AAVS1) safe harbor locus, with no detrimental effects^24–26^. Although ZFN and TALENs have shown great promise as targeted DNA editors, they are time consuming, expensive and challenging to engineer as new nuclease sequences must be generated for every new genomic target. Alternatively, clustered regularly interspaced short palindromic repeat/Cas9 (CRISPR/Cas9), which was developed by several groups in 2013 ^27–30^, allows for quicker, cheaper and easier-to-design human genome editing. CRISPR/Cas9 utilizes short guide RNAs (gRNAs; ∼20bp in length) to direct the Cas9 endonuclease to a specific genomic locus and induce a double stranded DNA break. Both the Cas9 enzyme and gRNA sequences can be encoded in a single plasmid and when co-transfected with a donor DNA plasmid can lead to higher homology directed repair (HDR) knock-in efficiency than previous editing tools^31^. We have previously described the first CRISPR-Cas9 system for AAVS1 integration of donor constructs containing an antibiotic resistance selection gene and both fluorescence (*tdTomato*) and bioluminescence (*Firefly luciferase*) reporter genes^32^. We were able to confirm the correct and stable integration of donor DNA at the AAVS1 site and functional reporter gene expression *in vivo*. However, some of the limitations of our study include (i) the low editing efficiency (∼3.8%) of HEK-293T cells; (ii) the use of large CRISPR/Cas9 and donor DNA plasmids that contained bacterial and antibiotic resistance genes – which limit transfection efficiency and would have associated safety concerns for clinical translation; and (iii) the lack of a translationally relevant reporter gene. In this study, we aimed to address these limitations by improving the efficiency and clinical safety of reporter gene integration at the AAVS1 safe harbor site and included a translationally relevant reporter gene.

We posited that the low editing efficiency of our first system was due in part to reduced transfection and knock-in efficiency, which is common with larger DNA plasmids, and the use of the HDR repair pathway for integration, which is intrinsically inefficient and not readily accessible to non-dividing cells^33^. In contrast to HDR-mediated DNA repair, the non-homologous end joining (NHEJ) pathway is active in both proliferating and non-proliferating cells and is generally considered more efficient than HDR in mammalian cells^34^. Recent studies have shown that by designing a CRISPR/Cas9 system that includes the same gRNA cut site in the donor vector as the genomic target site, the NHEJ repair pathway will more efficiently lead to transgene integration in zebrafish^35^ and mammalian cells^36,37^. Suzuki *et al.* refer to this mechanism as homology independent targeted integration (HITI), which is expected to lead to increased insertion in the forward rather than the reverse direction, as intact gRNA target sequences will be preserved in the latter^38^. Therefore, we postulated that HITI will increase the efficiency of reporter gene integration at the AAVS1 site (Figure 1A), compared to HDR. To address the problem of size and bacterial/antibiotic resistance genes in plasmids, our group and Suzuki *et al.*^38^, previously designed minicircles (MC) to express genes of interest^39,40^. First described by Darquet *et al.*^41^, MCs lack the bacterial backbone and antibiotic resistance genes that would otherwise compromise biosafety and clinical translation. In addition, the removal of the prokaryotic backbone also greatly reduces the size of the vector, thus improving transfection efficiency or providing space for the inclusion of other transgenes. To that end, we aimed to improve on our previous work by including a translationally relevant reporter gene in a multi-modality imaging HITI MC donor. We determined that the rat organic-anion-transporting polypeptide 1A1 (*Oatp1a1*) gene was an ideal candidate. *Oatp1a1* is a positive contrast magnetic resonance imaging (MRI) reporter gene due to its ability to uptake a clinically approved, liver-specific paramagnetic contrast agent called gadolinium ethoxybenzyl diethylenetriamine pentaacetic acid (Gd-EOB-DTPA; Primovist/Eovist)^42^. We have previously shown that *Oatp1a1* is a sensitive, quantitative, MRI reporter for 3-dimensional cancer cell distribution *in vivo*^43^. The purpose of this study was to develop HITI MC donor vectors for CRISPR/Cas9 editing of cells at the AAVS1 locus with three reporter genes to allow for multimodality, longitudinal *in vivo* monitoring of their fate following transplantation.

**Figure 1.**
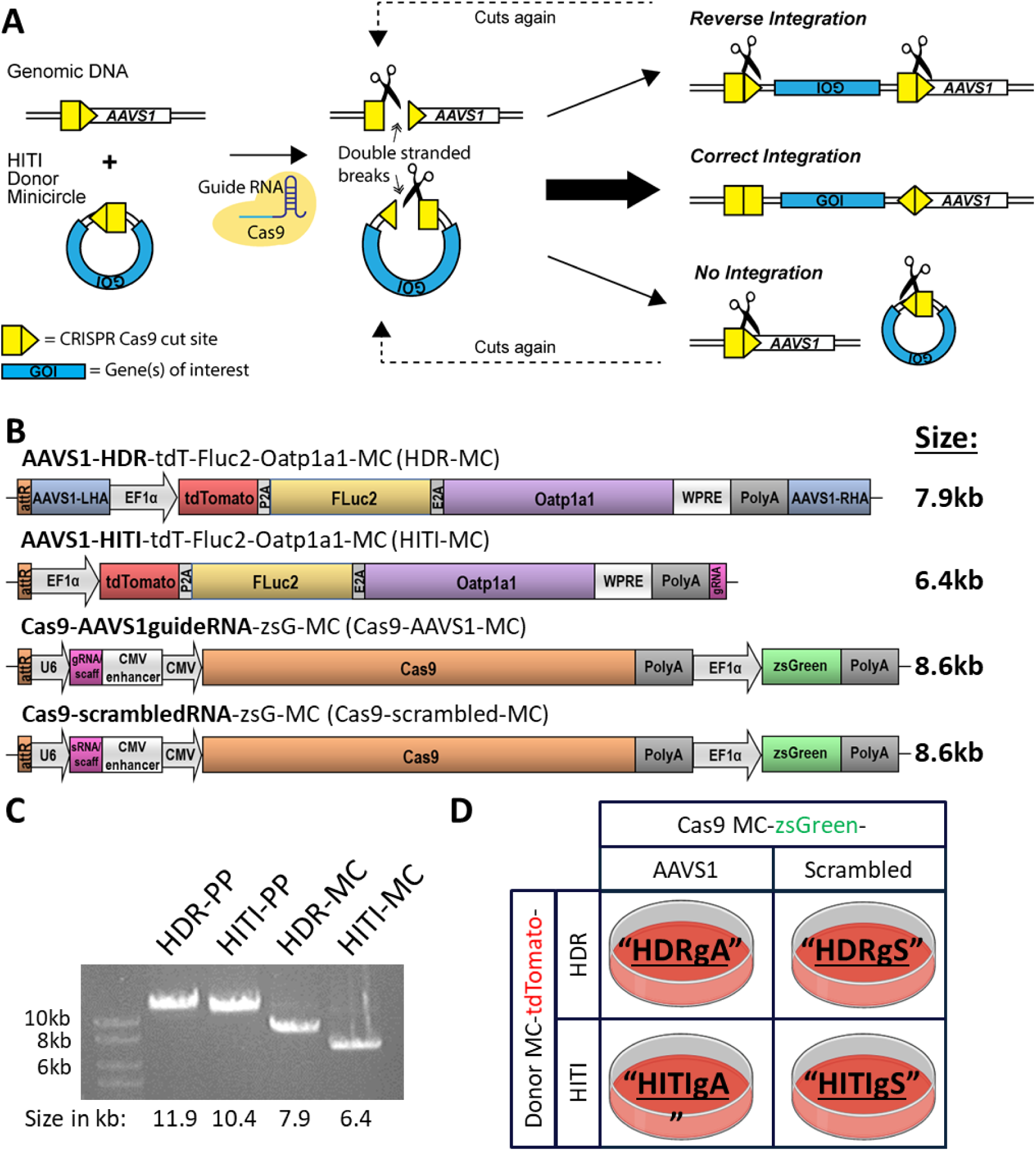
Homology-Independent Targeted Integration (HITI) experimental design. *A*, HITI minicircles (MCs) contain a Cas9 cut site identical to that at the AAVS1 safe harbor locus. Both genomic and MC DNA are cut in the presence of a guide RNA and Cas9. Genes of interest (GOI) are only stably integrated into the genome when inserted in the correct orientation, otherwise the Cas9 cut sites are preserved, which increases the likelihood of continuous Cas9 cutting. Image adapted from Suzuki *et al.*, 2016. *B*, Trimodality HDR, HITI and Cas9 MC constructs designed for this study. *C*, Restriction digest agarose gel of parental plasmids (PP) and MCs and indicated band sizes. *D*, transfection regimen for combinations of donor and Cas9 MCs and simplified abbreviations for each condition.

## Materials and Methods

### Constructs

Construct designs are shown in Figure 1B. The pCas9-AAVS1guideRNA-zsG-MC (Cas9-AAVS1-MC) and pCas9-scrambledRNA-zsG-MC (Cas9-scrambled-MC) parental plasmids originated from pCas-Guide-AAVS1 and pCas-Guide-Scrambled plasmids purchased from Origene (Maryland, USA). The Cas9 enzyme and guide RNA sequences were cloned between *attB* and *attP* recombination sites in a minicircle bacterial backbone containing a ZsGreen (*zsG*) fluorescence reporter driven by the elongation factor 1-α promoter (*hEF1α*). The AAVS1-HDR-tdT-Fluc2-Oatp1a1-MC (HDR-MC) construct was derived from an HDR vector lacking the *Oatp1a1* gene as we described previously^32^. This plasmid is driven by the *hEF1α* promoter, expresses tdTomato (*tdT*), firefly luciferase (*Fluc2*) and Organic anion transporting polypeptide 1a1 (*Oatp1a1*) using a self-cleaving 2a peptide system. For improved expression, the plasmids also contain the woodchuck hepatitis virus post-transcriptional regulatory element (*WPRE*) followed by the human growth hormone polyadenylation signal (*hGH* polyA). The HDR plasmid contains the left and right homologous arms (RHA:527bp, LHA:481bp) that are complementary to the region flanking the AAVS1 cut site; the homologous arms were obtained from the pAAVS1-puroDNR plasmid from Origene (Maryland, USA). The *Oatp1a1* gene was added through PCR amplification from a previously made vector we constructed using PGK_Straw_E2A_Oatp1a1 (a kind gift from Dr. Kevin Brindle’s laboratory; University of Cambridge). Using the HDR-MC parental plasmid as the template, we generated the pAAVS1-HITI-tdT-Fluc2-Oatp1a1-MC (HITI-MC) parental plasmid using the In-Fusion cloning kit from Clontech (Takara Bio, California, USA). Using restriction enzyme digestion, we extracted the bacterial backbone and minicircle recombination sites and then extracted the three reporter genes (without the homologous arms); *tdT, Fluc2* and *Oatp1a1* from the HDR-MC construct using PCR. However, for HITI functionality we designed our primers to also include a 23 bp extension (5’-GTTAATGTGGCTCTGGTTCTGGG-3’) downstream of the polyA sequence, which incorporates the same cut site and protospacer adjacent motif (PAM) sequence for our AAVS1 gRNA which allows for Cas9 cutting of both the MC and genomic DNA.

### Minicircle Production

ZYCY10P3S2T E. coli (System Biosciences, Palo Alto, CA, USA) were transformed with the original parental plasmids (PP) of all four constructs; HDR-MC or HITI-MC, Cas9-AAVS1-MC and Cas9-scrambled-MC and viable colonies selected using kanamycin plates. Colonies were picked 24 hrs after transformation, grown in 6 ml of lysogeny broth (LB) with kanamycin for 6 hrs at 37°C, followed by growth in terrific broth (TB) for 12 hrs at 37°C. To induce expression of the phiC31 integrase for MC production via *attB* and *attP* recombination, 100 ml of LB broth together with 100 µl of 20% arabinose induction solution (System Biosciences, Palo Alto, CA, USA) and 4 ml of 1N NaOH was added to the culture and grown for 5.5 hrs at 30°C. An endotoxin-free maxi kit (Qiagen, Valencia, CA, USA) was used to purify both PP and MC. Following purification of the MC products, PP contamination was removed using the Plasmid Safe ATP-dependent DNase kit (Epicentre, WI, USA), and the products where cleaned and concentrated using the Clean & Concentrator-25 kit (Zymo Research, CA, USA).

### Cell Culture and Transfection

Human embryonic kidney 293T cells and human adenocarcinoma HeLa cells (both from ATCC, Manassas, VA, USA) were grown in DMEM medium (Wisent Bioproducts, Québec, Canada) supplemented with 10% Fetal Bovine Serum (FBS; Wisent Bioproducts, Québec, Canada) and 1x Antibiotic-Antimycotic (ThermoFisher, Waltham, MA, USA). Human grade 4 adenocarcinoma PC3 cells were a kind gift from Dr. Hon Leong (Western University, ON, Canada) and were grown in RPMI (Wisent Bioproducts, Québec, Canada) supplemented with 5% FBS and 1x Antibiotic-Antimycotic. Cells were transfected with the linear polyethylenimine transfection agent jetPEI (Polyplus-transfection, Illkirch, France), according to the manufacturer’s instructions. Briefly, cells were grown in 6-well plates until 80-90% confluency and co-transfected with 1 µg each of Cas9-AAVS1-MC or Cas9-Scrambled-MC together with 1 µg of the donor MC constructs: HDR-MC or HITI-MC, for a total DNA mass of 2 µg. The DNA was prepared in 150 mM NaCl and complexed with 4 µl of jetPEI reagent per well.

### FACS and Flow Analysis

All FACS and flow cytometry was performed at the London Regional Flow Cytometry Facility (Robarts Research Institute, London, Canada). Forty-eight hours post transfection, the population of cells displaying both red (tdTomato) and green (zsGreen) fluorescence were sorted using a BD FACSAria III cell sorter (BD Biosciences, San Jose, CA, USA). At selected timepoints following FACS the cells were analyzed for tdTomato fluorescence using a FACSCanto flow cytometer (BD Biosciences, San Jose, CA, USA). Either 14- or 21-days post the initial sort, the cells were again sorted on the FACSAria III to purify tdTomato positive cells only (referred to as the pooled population). In this regard, our protocol aimed to sort cells that had incorporated the MC inserts (based on tdTomato fluorescence) into the genome and excluded any cells that had randomly integrated Cas9 MC DNA (zsGreen). At the same time as the second (tdTomato) sort, individual cells were plated into wells of a 96-well plate to enable single cell colonies to be grown and expanded (referred to as clonal cell populations).

### Genomic DNA Extractions and AAVS1 Integration Analysis

Extraction of genomic DNA from the pooled population of cells was performed using the DNeasy Blood and Tissue kit (Qiagen, Valencia, CA, USA) following manufacturer’s instructions. DNA quality and concentrations were measured on a NanoDrop 1000 spectrophotometer (ThermoFisher). Extraction of genomic DNA from clonal populations was performed as we described previously^32^. Briefly, cell pellets were resuspended in a QuickExtract DNA extraction solution (Lucigen, Middleton, WI, USA), incubated at 65°C for 10 mins, vortexed and incubated at 98°C for 5 mins. The DNA was then directly used for PCR or stored at −20°C. To check for integration at the AAVS1 site, two primers where designed to amplify the 3’ junction between the donor cassette and the AAVS1 site outside of the homologous arm region. The forward primer was uniquely complementary to the polyA tail in the MC cassette (5’-CCTGGAAGTTGCCACTCCAG-3’) and the reverse primer to the AAVS1 site (5’-AAGGCAGCCTGGTAGACAGG-3’). A 1.3 kb PCR product was produced if the MC-HDR was correctly integrated at the AAVS1 site and a 1.7 kb PCR product if MC-HITI was correctly integrated. GAPDH primers were designed as DNA loading controls and to confirm successful DNA extractions: forward 5’-TTGCCCTCAACGACCACTTT-3’ and reverse 5’-GTCCCTCCCCAGCAAGAATG-3’ and yielded a PCR product of 502 bp. Agarose gel electrophoresis with 1% agarose gels and RedSafe (FroggaBio, ON, Canada) was used to separate and visualize PCR products.

### *In Vitro* Fluorescence and Bioluminescence Imaging

The pooled and clonal cell populations were evaluated for tdTomato fluorescence expression on an EVOS FL auto 2 microscope (ThermoFisher, Waltham, MA, USA). For BLI experiments, varying cell numbers were plated in triplicates into black walled 96-well plates. D-luciferin (0.1 mg/ml; PerkinElmer, Waltham, MA, USA) was added to each well and images rapidly collected on an IVIS Lumina XRMS In Vivo Imaging System (PerkinElmer) equipped with a cooled CCD camera. Average radiance values in photons/sec/cm^2^/steradian were measured from regions of interest drawn around each well using LivingImage software (PerkinElmer).

### *In Vitro* Magnetic Resonance Imaging

Naïve and Oatp1a1-expressing cell clones (2×10^6^) were seeded in T-175 flasks and grown for 3 days. Cells were incubated with media containing 6.4 mM Gd-EOB-DTPA or with media containing an equivalent volume of PBS for 90 minutes at 37°C and 5% CO_2_. Cells were then washed 3 times with PBS, trypsinized and pelleted in 0.2 mL Eppendorf tubes, and placed into a 2% agarose phantom mould that was incubated in a 37°C chamber for two hours to mimic body temperature. MRI was performed on a 3-Tesla GE clinical MRI scanner (General Electric Healthcare Discovery MR750 3.0 T, Milwaukee, WI, USA) and a 3.5-cm diameter birdcage RF coil (Morris Instruments, Ottawa, ON, Canada). A fast spin echo inversion recovery (FSE-IR) pulse sequence was used with the following parameters: field of view (FOV) = 256 × 256, repetition time (TR) = 5000 msec, echo time (TE) = 19.1 msec, echo train length (ETL) = 4, number of excitations (NEX) = 1, receiver bandwidth (rBW) = 12.50 MHz, inversion times (TI) = 50, 100, 125, 150, 175, 200, 250, 350, 500, 750, 1000, 1500, 2000, 2500, 3000 msec, in-plane resolution = 0.27 mm^2^, slice thickness = 2.0 mm, scan time = 5 min and 25 sec per inversion time. Spin-lattice relaxation rates were computed via MatLab by overlaying the image series and calculating the signal intensity on a pixel-by-pixel basis across the inversion time image series, followed by a fitting of the data into the following equation to output the spin-lattice relaxation time, where S represents signal intensity, κ represents the scaling factor, and ρ represents proton spin density:

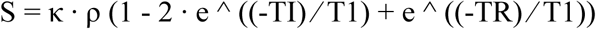

### Animal models

All animal protocols were approved by the University Council on Animal Care at the University of Western Ontario (Protocol #2015-058) and follow the Canadian Council on Animal Care (CCAC) and Ontario Ministry of Agricultural, Food and Rural Affairs (OMAFRA) guidelines. Crl:NU-*Foxn1*^*nu*^ (nude) male mice (Charles River Laboratories, Wilmington, MA, USA; N = 3-5) aged 6-8 weeks were used for subcutaneous tumour model injections and NOD.Cg-*Prkdc*^*scid*^ *Il2rg*^*tm1WjI*^/SzJ (NSG) immunodeficient male mice (obtained from the Humanized Mouse and Xenotransplantation Facility at the Robarts Research Institute, University of Western Ontario, London, Canada; N = 3) for experimental metastasis models (intravenous cell injections).

### *In Vivo* Bioluminescence Imaging

BLI was performed on the same IVIS Lumina XRMS system described for *in vitro* imaging. Mice were anesthetized with 2% isoflurane in 100% oxygen using a nose cone attached to an activated carbon charcoal filter for passive scavenging and kept warm on a heated stage. BLI images were acquired with automatic exposure times until the peak BLI signal was obtained (up to 40 mins). Regions of interest were manually drawn using LivingImage software to measure average radiance (photons/sec/cm^2^/sr). The peak average radiance was used for quantification for each mouse.

### *In Vivo* Magnetic Resonance Imaging and Quantification

All mouse MRI scans were performed with a custom-built gradient insert and a bespoke 5 cm diameter solenoidal RF coil, as we described previously^43^. Mice were kept anaesthetized during the scan with 2% isoflurane administered via a nose cone attached to the coil. *T*_1_-weighted images were acquired using a spoiled gradient recalled acquisition in steady-state pulse sequence using the following parameters: field of view, 50 mm; repetition time, 14.7 msec; echo time, 10.5 msec; receiver bandwidth, 31.25 MHz; echo train length, 4; frequency and phase, 250 × 250; flip angle, 60 degrees; number of excitations, 3; 200 μm isotropic voxels; scan time, approximately 15 minutes per mouse. Pre-contrast images were acquired followed by administration of 1.67 mmol/kg Gd-EOB-DTPA (Primovist; Bayer, Mississauga, ON, Canada) via the tail vein. Mice were then re-imaged 20 minutes later for immediate post-contrast images, which provide positive contrast to many tissues, including the naïve tumours, as a result of Gd-EOB-DTPA pooling; and/or 5 hours later for Oatp1a1-specific uptake. This time-point was determined to allow enough time for Gd-EOB-DTPA to be cleared, yet still provide strong positive contrast in Oatp1a1-expressing cells^42,43^. Contrast-to-noise ratio (CNR) measurements were calculated from MR images using ITK-snap open source software (www.itksnap.org)^44^. Tumours were manually segmented in 3 dimensions by tracing the tumour or control tissue (hind leg muscle) with polygon and paintbrush tools and pixel intensity recorded in every slice. The CNR of tumours was calculated by taking the signal intensity of the difference between tumour regions and muscle tissue divided by the standard deviation of background signal

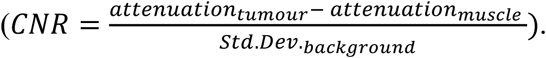

### Oatp1a1-induced Gd-EOB-DTPA uptake MRI and BLI sensitivity

To evaluate the cellular detection sensitivity of Oatp1a1 expressing cells with Gd-EOB-DTPA-enhanced MRI, nude mice were injected with 50 µl of cell suspensions in PBS containing 3×10^6^ total cells/injection at the following ratios: 3×10^6^ naïve cells alone; 10^4^ PC3-HITI + 2.99×10^6^ naïve cells; 10^5^ PC3-HITI + 2.9×10^6^ naïve cells; 10^6^ PC3-HITI + 2×10^6^ naïve cells; and 3×10^6^ PC3-HITI cells alone, subcutaneously in 5 locations on the back/flank region. Immediately after cell injections, 1.67 mmol/kg Gd-EOB-DTPA was injected into the tail vein and mice were imaged on a 3T clinical grade MR scanner 5 hours later. This time point allows for clearance of Gd-EOB-DTPA from the body yet provides sufficient time for the agent to penetrate the subcutaneous injections sites and accumulate in Oatp1a1 expressing cells. After MRI, mice were moved to the IVIS scanner and injected with 100 µl of 30 mg/ml D-Luciferin intraperitoneally and BLI was performed, as described earlier.

### 293T and PC3 tumour models

293T or PC3 naïve and HITI engineered cells were injected subcutaneously (2.5×10^6^ 293Ts and 1×10^6^ PC3s) on the left and right flanks of nude mice, respectively (293T, N = 2; PC3, N = 5). For experimental metastasis studies, 5×10^5^ PC3 naïve or HITI engineered cells were injected into the tail veins of NSG mice (N = 3). Tumour growth was tracked on a weekly basis with BLI, as described above. MRI was performed on mice at various timepoints, as indicated in the results section. Firstly, a pre-contrast scan was performed on all mice, followed immediately with injection of the Gd-EOB-DTPA contrast agent into the tail vein (1.67 mmol/kg). For some experiments the mice were re-scanned 15-20 mins after contrast injection to show tumour and whole-body distribution of Gd-EOB-DTPA. In all instances MRI scans were performed ∼5 hours post-contrast injection since Oatp1a1 expressing cells still retain Gd-EOB-DTPA and show strong positive contrast at this time point. This also allows enough time for washout of Gd-EOB-DTPA in most tissues and organs (except for the gastrointestinal tract and bladder where cleared Gd-EOB-DTPA accumulates before being excreted)^42^.

### Statistics

Statistical analysis was performed with GraphPad Prism version 7 (GraphPad Software Inc., CA, USA; www.graphpad.com) software. One-way ANOVA with Tukey’s multiple comparison test was used for *in vitro* and *in vivo* BLI and CNR data analysis. An unpaired one-tailed *t*-test was used to analyze the increase in CNR for PC3-HITI day 11 vs day 46 tumours.

## Results

### CRISPR/Cas9 engineering of multiple human cell types with tri-modal reporter gene minicircle donors

In this study, we designed our trimodal reporter gene system in MC constructs to reduce the size and immunogenicity of our donor DNA and to remove antibiotic resistance genes. To compare the efficiency of HDR vs HITI editing at the AAVS1 site, we designed two donor and two Cas9-expressing MCs, as shown in Figure 1B. The HDR and HITI constructs were engineered to express tdTomato (*tdT*), firefly luciferase (*Fluc2*) and rat organic anion transporting polypeptide 1a1 (*Oatp1a1*) genes under the control of an *EF1α* promoter and 2A self cleaving peptide system (Figure 1B). The HDR and HITI parental plasmids initially measured 11.9 kb and 10.4 kb in size, which was then reduced to 7.9 kb and 6.4 kb when recombined into MCs, respectively, as confirmed by agarose gel electrophoresis (Figure 1C). The HDR-MC was flanked by left and right AAVS1 homologous arms either side of the AAVS1 genomic cut site, whereas the HITI donor contained the same CRISPR/Cas9 cut site as the AAVS1 genomic site (Supp. Figure 1). In this instance, if the MC DNA inserted into the correct orientation at the AAVS1 site, the CRISPR/Cas9 cut sites would be lost and the trimodal reporter genes would be stably integrated into the genome (Supp. Figure 1). The Cas9 expressing MCs were designed to contain the necessary RNA scaffolding and gRNA sequences targeting the AAVS1 site or a scrambled gRNA control, alongside a zsGreen (*zsG*) fluorescent reporter gene (Figure 1B). Both the pCas9-AAVS1-MC and pCas9-scrambled-MC constructs measured 12.5kb in parental plasmid form, and 8.6kb in MC form (Figure 1B).

Our first objective was to determine the correct integration of our donor MCs in three human cell lines; HEK 293T, HeLa and PC3 cells. All 3 cell lines where co-transfected with the HDR-MC or HITI-MC together with either the Cas9-AAVS1-MC or Cas9-scrambled-MC (as outlined in Figure 1D) and grown for 48hrs. The cells were then FACs sorted for tdT+/zsG+ cells in order to purify cells that were successfully co-transfected, and tdT fluorescence was then tracked every 7 days using flow cytometry (Supp.Figure 2A-B). In two separate experimental groups, the cells were then resorted 14 or 21 days later for tdT+/zsG- cells, to ensure that the cell populations had not randomly integrated the Cas9-zsG MCs into the genome (Supp. Figure 2C). Both PC3 experimental groups were resorted 14 days after the initial sort (and not 21 days later) due to lower transfection rates. However, resorting the cells 14 or 21 days later had a negligible effect on tdT+ cell populations across the timepoints. For almost all cell types there was a higher percentage of tdT+ fluorescence cells at endpoint in the HITI-AAVS1 groups (pink shading, Supp. Figure 2C), suggesting better or more stable integration compared to HDR-AAVS1 groups.

**Figure 2.**
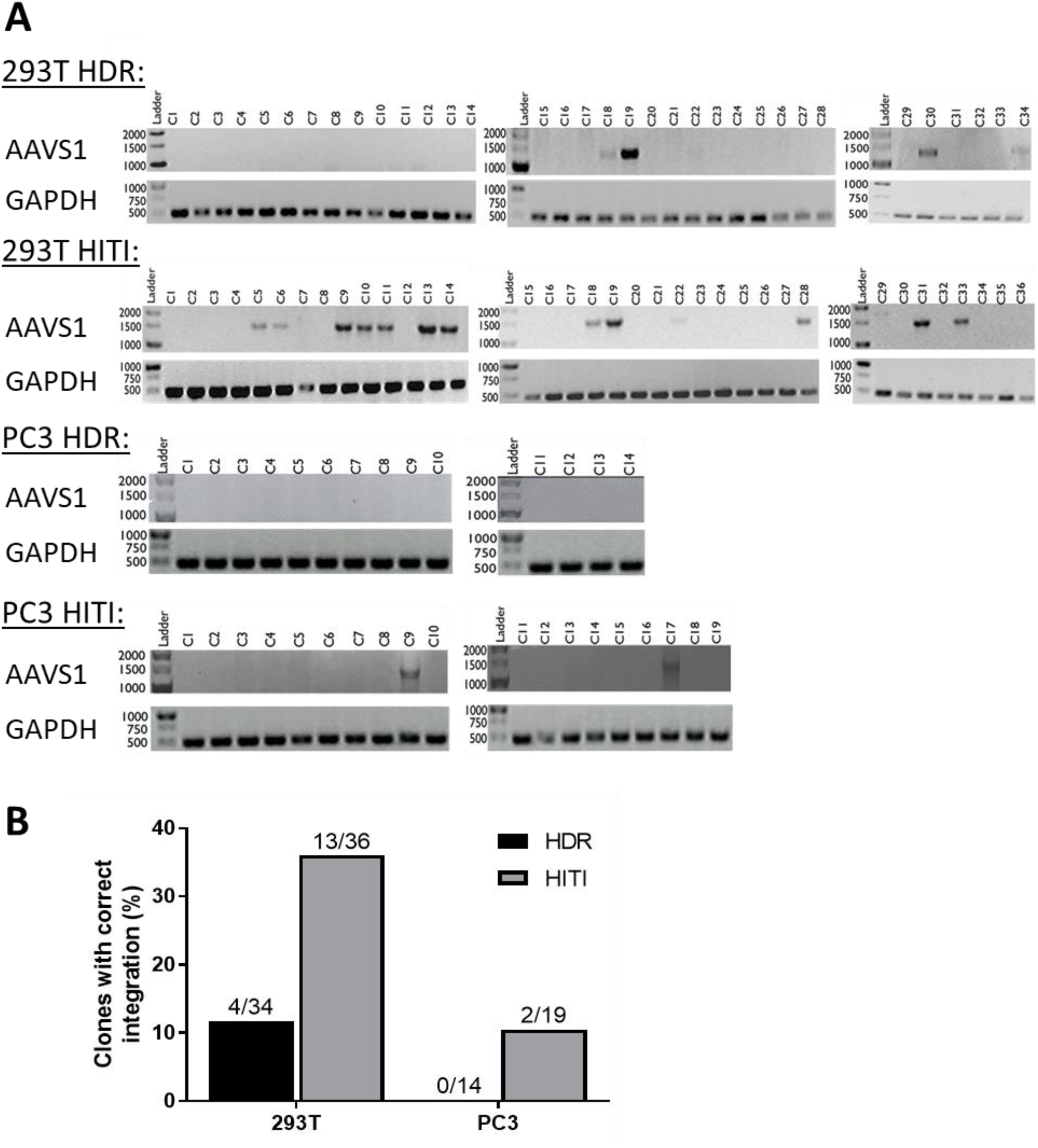
Junctional PCR integration checks for 293T and PC3 clonal cell populations. *A*, PCR integration checks at the AAVS1 site. GAPDH was amplified as a DNA loading control. *B*, Quantification shows a higher number of positive integration clones for HITI engineered cells compared to HDR for both 293T and PC3 cell lines.

### Mixed cell population (MCP) integration and BLI analysis

We next performed junctional PCR analysis on extracted DNA samples to determine whether the tdT+ MCPs had correctly incorporated the large, trimodal donor MCs into the AAVS1 site in the right orientation (Supp. Figure 3A). A correct integration band (1.4 kb) was detected for all HITI-guideAAVS1 (HITIgA) engineered cells (very low transfection efficiency for PC3 cells may explain why the integration band was weak) as well as a correct integration band (1.3 kb) for HDR-guideAAVS1 (HDRgA) cells for 293T and HeLa MCPs. There were no integration bands for the control naïve cells or cells engineered with scramble guide RNA (HITI/HDRgS). Next, we performed *in vitro* BLI experiments to determine if the integrated reporter gene was functioning in the MCPs. Varying numbers of each cell type were imaged with BLI after addition of D-luciferin to visualize *FLuc2* expression (Suppl Figure 3B). In all cell types, there was a positive correlation between BLI signal and cell number and, interestingly, a consistently higher signal was seen in all cell types engineered with HITI-guideAAVS1 compared with HDR-guideAAVS1.

**Figure 3.**
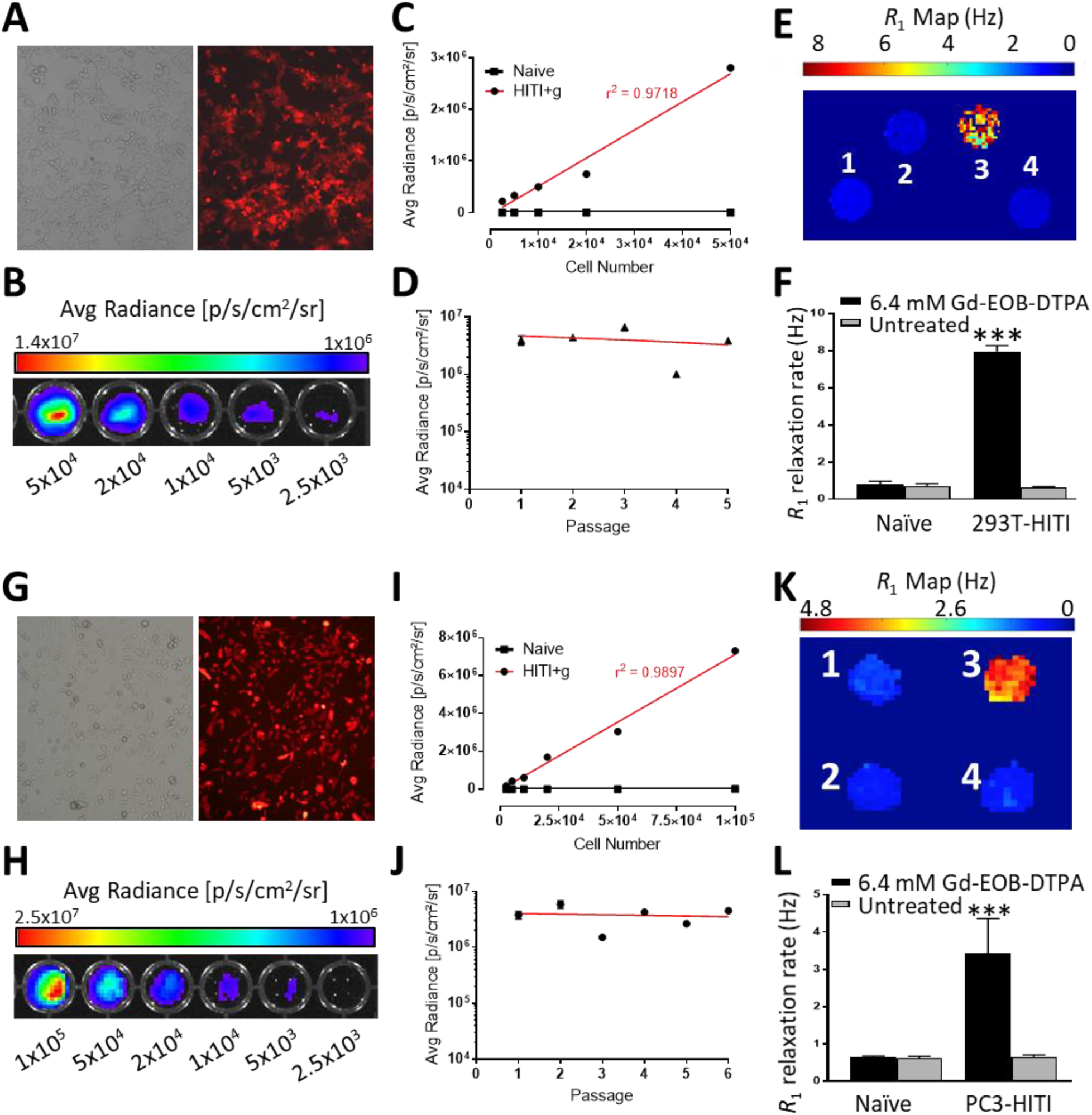
*In vitro* FLI, BLI and MRI characterization of 293T-HITI (*A-F*) and PC3-HITI (*G-L*) clones. *A* and *G*, brightfield and tdT fluorescence. *B* and *H*, BLI intensity maps related to cell number. *C* and *I*, quantification of BLI signal to cell number. *D* and *J*, BLI signal over successive passages. *E* and *K*, Spin-lattice relaxation maps of representative phantoms containing pellets of cells untreated or treated with 6.4 mM Gd-EOB-DTPA, as follows: 1, naive, treated; 2, naïve untreated; 3, HITI treated; 4, HITI untreated. *F* and *L*, quantification of spin-lattice rates (n = 3, *** *P* < 0.001).

### HITI is more efficient than HDR in clonal populations

Next, we used clonal cell isolation to determine whether HITI or HDR was more efficient at correctly integrating our large donor MCs at the AVVS1 site. Single cell tdT+ clones were isolated from the 293T and PC3 MCPs into 96-well plates during a third FAC sort. We decided to use the 293T cells as a proof-of-principle cell line and the PC3 cells as a relevant prostate cancer model cell line, hence the HeLa cells were not included in studies from this point onwards. PCR integration checks were performed on the 293T and PC3 clonal populations to determine the efficiency of HITI-vs HDR-mediated reporter gene integration at the AAVS1 site (Figure 2A-B). The number of 293T clonal populations with correct integration was 11.8% (4/34) for HDR-AAVS1 engineered cells and 36.1% (13/36) for HITI-AAVS1 clones (Figure 2A-B). PC3 cells grew fewer colonies but showed zero integration at the AAVS1 site for tdT+ HDR engineered cells (0/14), whereas 10.5% (2/19) of the HITI engineered colonies had correct reporter gene integration, indicating that HITI was more efficient in both cell types.

### *In vitro* reporter gene imaging

Next, we expanded single 293T and PC3 clonal populations that had correct integration bands for further *in vitro* reporter gene functionality testing. Firstly, we confirmed tdT fluorescence for both the 293T-HITI (Figure 3A) and PC3-HITI (Figure 3G) clones via fluorescence microscopy. In addition, there was a positive correlation between BLI signal and increasing cell number for 293T (Figure 3B-C; *r*^*2*^=0.9718) and PC3 (Figure 3H-I; *r*^*2*^=0.9897) cells. BLI signal measured over several passages showed stable FLuc2 expression over time for both clonal cell lines (Figure 3D and J). To test for Oatp1a1 functionality, 293T naïve, 293-HITI, PC3 naïve and PC3-HITI cells were incubated with or without Gd-EOB-DTPA (6.4 mM) in normal media for 90 minutes, washed thoroughly, and pelleted before inserting into an agarose phantom. Inversion recovery MRI was performed at 3 Tesla and 37°C, and spin-lattice relaxation rate (*R*_1_) maps were generated (Figure 3E, K). Neither the naïve 293T/PC3 or untreated 293T-HITI and PC3-HITI cell populations exhibited any change in *R*_1_ rates (Figure 3F, L). Only HITI clones expressing Oatp1a1 had significantly increased *R*_1_ rates after Gd-EOB-DTPA incubation, with ∼10-fold increase for 293T-HITI cells (7.952 ± 0.87 s^-1^) compared with naïve, treated controls (0.806 ± 0.038 s^-1^; n = 3, *P* < 0.001; Figure 3F) and ∼5-fold increase for PC3-HITI cells (3.426 ± 0.217 s^-1^) compared with naïve, treated controls (0.6402 ± 0.045 s^-1^; n = 3, *P* < 0.001; Figure 3L).

### Oatp1a1 sensitivity *in vivo*

To investigate the MR detection limit of Oatp1a1-expressing cells we injected various combinations of PC3-naïve and PC3-HITI cells at five sites subcutaneously on the backs of nude mice (Figure 4 and Supp. Figure 4). A total of 3×10^6^ cells were injected per site with the following number of PC3-HITI cells: 1 – 0 (naïve cell only control); 2 – 10^4^; 3 – 10^5^; 4 – 10^6^; 5 – 3×10^6^ (HITI cell only control). Naïve cells were included with HITI cells so that each injection contained a total of 3×10^6^/site. BLI signal intensity increased as PC3-HITI cell numbers increased (representative mouse shown in Figure 4A), with 10^6^ and 3×10^6^ HITI injections showing significant signal increase above naïve background controls (Figure 4B). Transverse MR images from the same mouse showed positive contrast at both the 10^6^ and 3×10^6^ HITI injection sites 5 hours after Gd-EOB-DTPA injection (Figure 4C). Similar to the BLI data, these sites also exhibited significantly higher contrast-to-noise ratios (CNR) than naïve controls (Figure 4D). The 10^4^ and 10^5^ PC3-HITI injections were difficult to visualize on MRI, had no discernible positive contrast and were, therefore, not measured. These data were consistent across all three mice (see Supp. Figure 4) and showed that the very minimum number of Oatp1a1-expressing cells we could detect with Gd-EOB-DTPA-based MR contrast was 10^6^ cells in a 50 μl injection volume.

**Figure 4.**
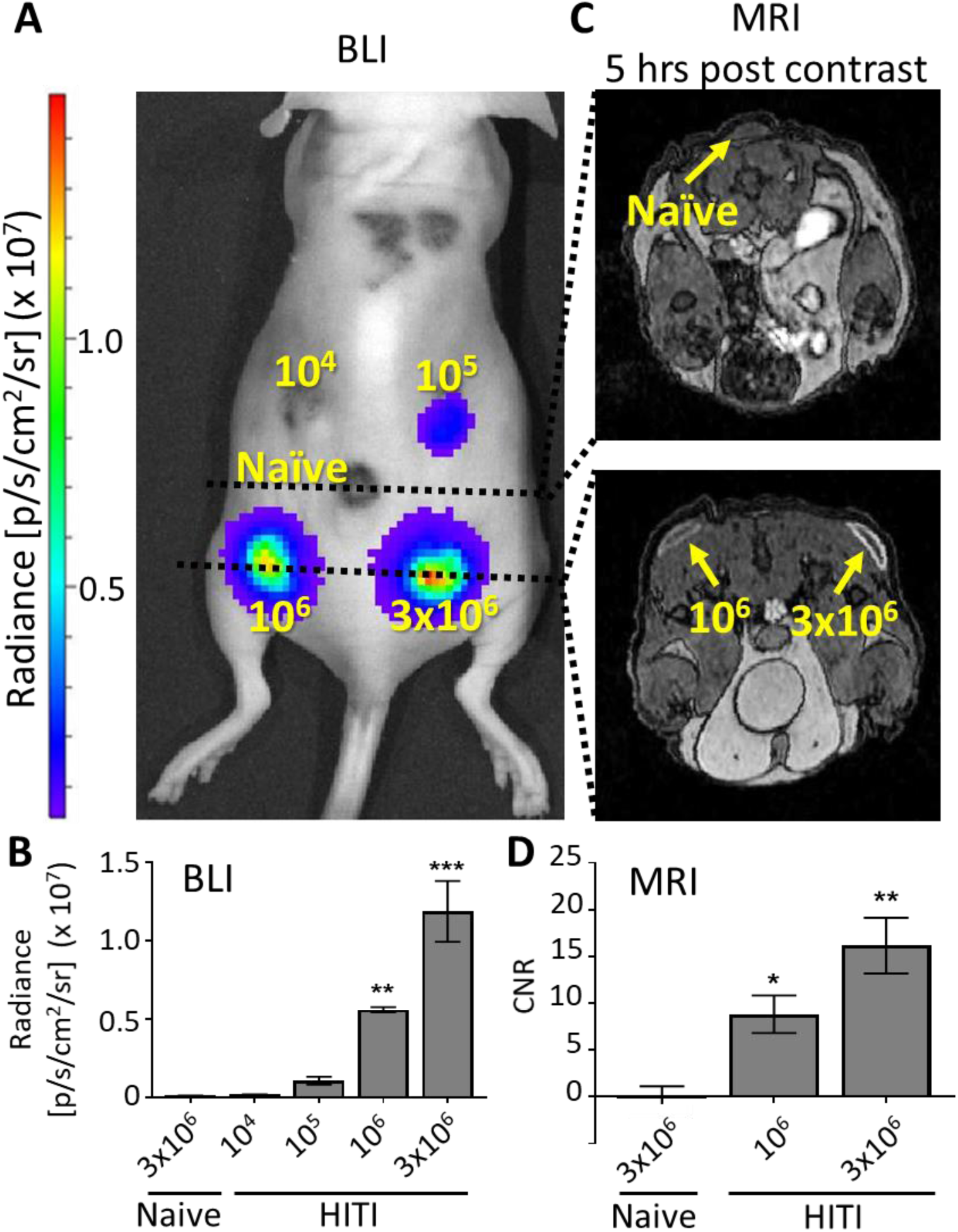
BLI and Oatp1a1 sensitivity *in vivo. A*, A total of 3×10^6^ PC3 cells were injected subcutaneously into five locations on the back of nude mice, with increasing concentrations of HITI engineered cells, as indicated in yellow, and corresponding BLI signals. *B*, quantification of BLI signal from ROIs around the five PC3 injection sites. *C*, MRI transverse views of cell injection sites 5 hours after Gd-EOB-DTPA injection. *D*, quantification of contrast-to-noise (CNR) ratio for 10^6^ and 3×10^6^ PC3-HITI cells. Note, 10^4^ and 10^5^ PC3-HITI injections lacked enough contrast to measure CNR values. n = 3 mice, * *P* < 0.05, ** *P* < 0.01, *** *P* < 0.001.

### PC3-HITI Oatp1a1 tumour models for MRI detection

As a proof-of-principle that our HITI-engineered cells could show Gd-EOB-DTPA induced positive MRI contrast in subcutaneous tumours, we injected 293T-naïve and 293T-HITI cells on either flank of a nude mouse (Supp. Figure 5). For both cell types, the large masses were visible on pre-contrast images but also showed noticeable positive contrast 20 minutes post-Gd-EOB-DTPA injection. However, 5 hours post-contrast, the naïve tumour had returned to pre-contrast background levels, whereas the HITI tumour had very prominent positive contrast that also showed heterogeneity within the tumour mass (Supp. Figure 5), similarly to what we reported previously^43^.

**Figure 5.**
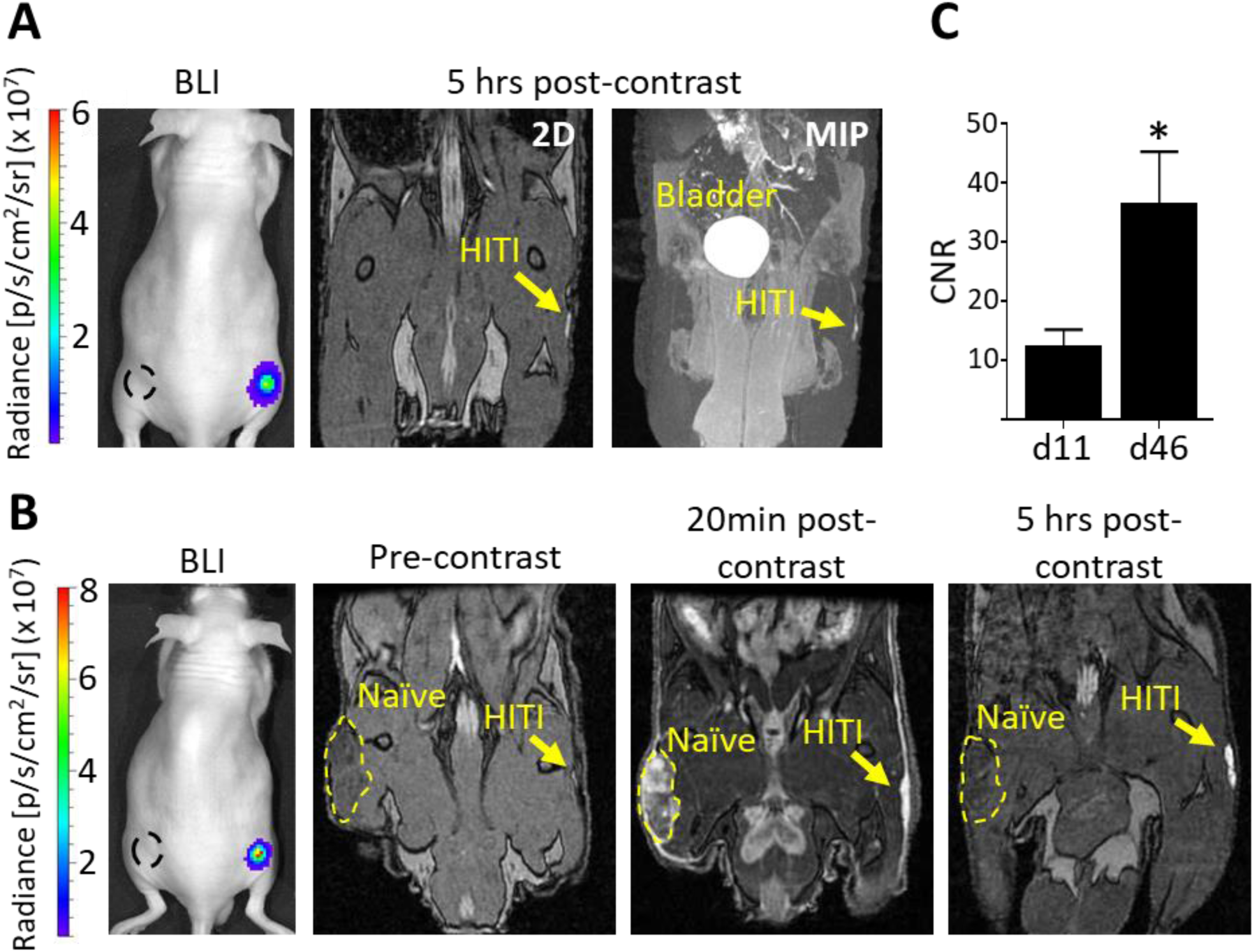
Longitudinal *in vivo* MRI of subcutaneous PC3-HITI cells. Mice were injected subcutaneously with 1×10^6^ naïve and PC3-HITI cells on the left and right flanks, respectively. BLI signal was present on right flank only. Naïve tumour locations are denoted by black dashed line. *A*, day 11 post PC3 injection. 2D and maximum intensity projection (MIP) images acquired 5 hrs post Gd-EOB-DTPA injection. *B*, the same mouse was re-imaged at day 46. Pre-, 20min post- and 5hr post-contrast coronal images were obtained. *C*, contrast-to-noise ratios (CNR) of PC3-HITI tumours 5hr post-contrast showed significant increase from day 11 to day 46. n = 4 (day 11) and n = 3 (day 46) * *P* < 0.05.

Moving into a more relevant cancer model, we next injected PC3-naïve and PC3-HITI clonal cells subcutaneously on either flank of nude mice and followed BLI development over time (Figure 5). After only 11 days post-injection, before the tumours were visible or palpable, clear positive contrast was observed for HITI engineered cells 5 hours after Gd-EOB-DTPA injection (Figure 5A and Supp. Figure 6). The same mice were then imaged at day 46 where the naïve tumour was visible due to pooled Gd-EOB-DTPA after 20 minutes post-contrast injection. In a similar fashion 5 hours post contrast, only the HITI engineered cells retained the Gd agent and showed bright, positive contrast (Figure 5B and Supp. Figure 7). Quantification of CNR over time showed a significant increase for the HITI tumours (Figure 5C). These data suggest that the Oatp1a1 MRI reporter can detect tumour burden at stages where the tumours are not visible or palpable and tumour growth can be tracked longitudinally with Gd-EOB-DTPA-enhanced MRI.

**Figure 6.**
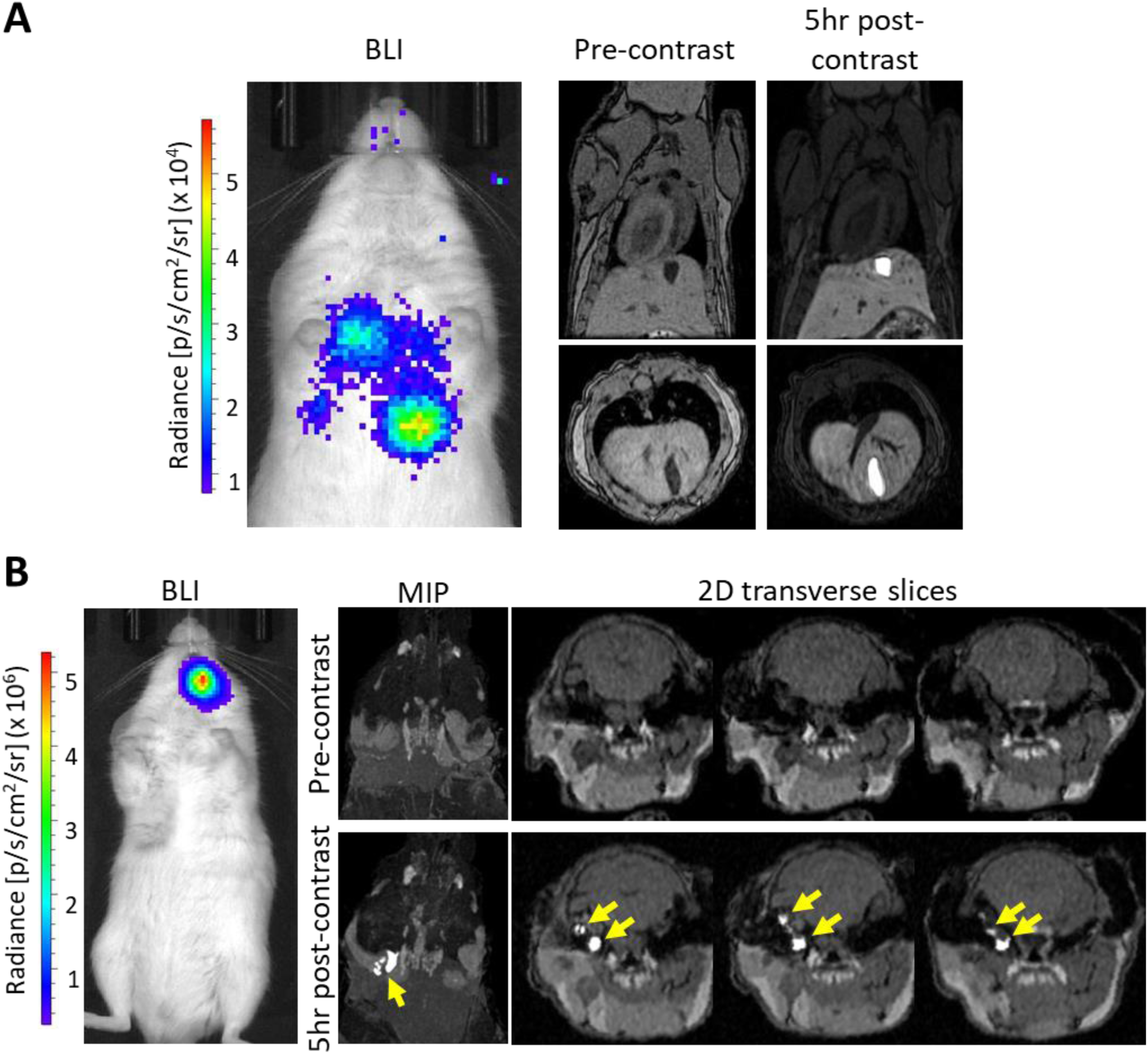
*In vivo* MRI detection of metastatic PC3-HITI tumours. *A*, The brightest BLI signal corresponded to a tumour in the liver, which is evident as a shadow in pre-contrast images and as a bright positive Gd-EOB-DTPA contrast tumour in images taken 5 hrs post-contrast injection. *B*, A mouse showed BLI signal in the head region, which was not evident in pre-contrast MIP and 2D transverse MRI slices (upper panel). However, clusters of PC3-HITI tumours were easily discernible in 5hr post-contrast images (yellow arrows, lower panel).

Finally, PC3-HITI cells were injected via the tail vein into NSG mice to investigate the ability of Oatp1a1 as a reporter gene for visualizing metastases (Figure 6). Using BLI as a guide, we were able to detect a Gd-EOB-DTPA-enhanced metastatic tumour in the liver of one mouse (Figure 6A). In this case, it was evident that a shadow in the liver (as seen in the pre-contrast MR image) was likely the tumour, which the post-contrast image confirmed. However, in another mouse that had BLI signal in the head region, a cluster of smaller PC3-HITI tumours was only identifiable in post-contrast images (Figure 6B), indicating the usefulness of this reporter gene for detecting metastatic burden with contrast-enhanced MRI.

## Discussion

As personalized medicine and CRISPR-editing become a reality in the clinic, there is a greater need to 1) improve the efficiency, efficacy and safety of genetically engineered cell-therapies, and 2) improve our understanding of disease progression and treatment response in preclinical models of disease. Reporter gene-based imaging allows us to track the location, viability, growth and efficacy of such treatments, and in preclinical models of cancer progression and treatment. In this study, we have developed a non-viral vector-based engineering system for large DNA multimodality reporter gene integration into the AAVS1 safe-harbour genomic locus. To improve safety further, we utilized MCs as the DNA vector of choice, which eliminates bacterial DNA contamination and antibiotic resistance genes. In addition, we showed that utilizing the non-homologous end joining (NHEJ) repair pathway with HITI could improve DNA editing efficiency in human cells compared to the more commonly used HDR pathway. Finally, building off our previous work^32^, we have engineered a trimodality reporter gene construct that contains a clinically relevant MRI reporter, *Oatp1a1*, in addition to fluorescent and bioluminescent genes which enabled cell sorting and non-invasive BLI/MRI of engineered cells in a pre-clinical cancer model.

One of the major limitations of engineering cells with large, multimodality reporter gene DNA plasmids is the reduced efficiency of both transfection and gene editing with increasing construct/insert size^45–47^. In addition, the presence of bacterial and antibiotic resistance genes in parental plasmids has the potential to exert immunological responses and raises safety concerns. To circumvent these issues, we designed our study to use MCs, which removes the bacterial backbone from parental plasmids and thus reduces the size of the DNA donor constructs. Using MCs instead of parental plasmids allowed us to remove ∼4 kb of unwanted DNA from our HDR construct, with a further reduction of ∼1.5 kb for the HITI MC when the homologous arm sequences were replaced with a 20 bp gRNA sequence (saving a total of ∼5.5 kb). These large-scale reductions thus provided us with room to upgrade our dual-modality *tdTomato* and *FLuc2* reporter gene construct we previously reported^32^ to a trimodality reporter gene construct with the addition of the *Oatp1a1* MRI reporter^42,43^. To improve safety and translatability we also removed the puromycin resistance gene to reduce the MC size by a further 600 bp and utilized FACS of tdTomato positive cells to obtain mixed and clonal cell populations instead of antibiotic selection. Our final step for improving safety was to design our system to target a “safe harbour” locus in genomic DNA. Several of these loci have now been reported in the literature^48^ and are described as sites where inserted genetic elements can function as intended, without causing alterations that would pose a risk to the host cell or organism^23^. For this study we targeted the AAVS1 site found within the human Protein Phosphatase 1 Regulatory Subunit 12C gene as this has been one of the best characterized, to date. No known side effects are associated with disrupting the PP1R12C gene, however it has been reported that mechanisms such as DNA methylation can silence transgenes targeted to this genomic region^49^. Since our studies rely on stable reporter gene expression over time for accurate cell detection and proliferation, we investigated whether reporter gene expression in our AAVS1 engineered 293T-HITI and PC3-HITI cell populations changed over time. We found that BLI signal was stable over several passages and tdT fluorescence was consistently expressed in both cell lines, indicating consistent transgene expression.

We have shown here that HITI-based CRISPR/Cas9 cell engineering is more efficient than the more commonly used HDR method for integrating large DNA donor constructs into the genome for stable expression. Targeted transgene integration is typically achieved using homologous arms and the HDR pathway, however, this mechanism is highly inefficient and is not usually active in non-dividing cells^33^. Indeed, our previous study showed only 3.8% of selected cells were correctly edited using the HDR mechanism^32^. In contrast, the HITI method which utilizes the NHEJ pathway is active in all stages of the cell cycle and in quiescent cells^36^, and thus has been used to improve editing efficiency. Using the method described by Suzuki *et al*.^38^, our engineered 293T and PC3 clonal cell populations did indeed have greater DNA integration at the AAVS1 site compared with HDR (36% and 12% for HITI vs 10.5% and 0% for HDR, respectively). However, it is important to note that the NHEJ repair pathway is error prone and often leads to insertions and deletions (indels) at the DNA junctions. Consequently, this mechanism is often taken advantage of to produce DNA disruptions, gene silencing and knock-outs. These issues would need to be considered if using the HITI method for correctional DNA editing and promoter-less vector integration, since these require specific DNA sequences, either upstream or downstream, to be preserved. In this case, we engineered cells with the only requirements being that the transgene inserts into the AAVS1 site (confirmed with junctional PCR) and that the reporter genes are consistently expressed (confirmed with imaging). Therefore, indels at either the 5’ or 3’ junction would likely have a negligible impact on our experiments.

Although we confirmed correct transgene integration at the AAVS1 site in our study, we cannot rule out integration at other off-target sites in HITI and HDR engineered populations. Several 293T and PC3 single-cell clonal populations expressed the *tdTomato* fluorescence reporter gene but did not show integration bands for the AAVS1 site. Evidence suggests that CRISPR/Cas9 is not 100% accurate and off-target effects have been reported^50,51^, thus, it is possible that the MCs integrated into off-target Cas9 cut sites. However, it is also plausible, and probably more likely, that the MCs inserted into the AAVS1 site in the wrong direction. Even though HITI is designed to minimize integration in the wrong orientation, the error-prone NHEJ repair mechanism of blunt-ended DNA breaks could lead to indels at the CRISPR/Cas9 cut site boundaries, which could then disrupt the ability of Cas9 to recognize and re-cut those sites. The likelihood of indel formation using Cas9 HITI could be reduced in future studies by adopting a similar method to that recently reported by Li and colleagues^52^. In that study, the authors utilize Cas12a, which leaves 5 bp overhangs, to precisely edit the genome in a process coined microhomology-dependent targeted integration (MITI)^52^. Independently of CRISPR, MCs, like plasmids, can also randomly integrate into the genome of cells, albeit at very low rates. Future work will need to analyze the rate of off-target integrations and possible indel disruptions at the CRISPR/Cas9 cut sites using techniques, such as next generation sequencing, to determine the full safety profile of HITI at safe harbour loci. To improve targeting specificity, studies have shown that high-fidelity Cas9 enzymes in ribonucleoprotein complexes (RNPs), instead of Cas9 DNA vectors, improve on-target activity, while reducing off-target editing^53,54^. In combination with RNPs, adeno-associated viruses (AAVs) are now commonly used as DNA donors for CRISPR experiments due to their high transduction capabilities in hard-to-transfect cell lines, their low risk of random integration and reduced immunogenic response. However, AAVs are still limited by their loading capacity of ∼4.5 kb, which would be a problem for large, multimodality imaging vectors as presented here, but conceivable for future studies where only one imaging reporter gene is required. With these emerging technologies, it is likely that CRISPR gene editing will become highly specific and thus safer in the near future.

We engineered cells with a multimodality reporter gene construct to enable us to go from single cell, optical imaging methods (FLI) to higher sensitivity whole-animal planar imaging (BLI) and superior 3D high-resolution tomographic imaging (MRI) in animals. This offers several advantages. Firstly, fluorescently activated cell sorting of tdTomato-expressing cells improves upon our previous study by eliminating the need for an antibiotic resistance selection gene, which constitutes a safety risk and has been associated with structural plasmid instabilities^55^. Secondly, the firefly luciferase gene (*FLuc2*), in combination with its substrate D-luciferin, allowed us to directly visualize engineered cells *in vivo* using BLI. Inclusion of bioluminescent genes in preclinical cancer models is a relatively inexpensive and valuable tool that also allows one to track cell migration and cell seeding in metastatic cancer models, assess cell viability and follow cell/tumour growth longitudinally^14^. A limitation of BLI is that it is restricted to small animal models of disease. However, it is useful for determining sites of cell arrest/seeding/growth and thus can be used in conjunction with other reporter genes as a guide for determining when and where to perform relatively more expensive, higher resolution clinical imaging, such as MRI^13^. To build off our previous dual FLI-BLI study^32^, we decided to include the MRI reporter gene, *Oatp1a1*, as a translationally relevant and sensitive reporter gene to complete our trimodality construct for HITI-based CRISPR engineering. First described by Patrick *et al.*^42^, *Oatp1a1* selectively, but reversibly, uptakes the clinically approved Gd^3+^ contrast agent Gd-EOB-DTPA and thus provides positive contrast in *T*_*1*_-weighted MR images. The authors concluded, therefore, that *Oatp1a1* engineered cells and tumours should be easier to detect than the negative contrast generated by *T*_*2*_-agents, such as such as superparamagnetic iron oxide (SPIO) and ferromagnetic agents^42,56^. In addition, engineering cells with integrated *Oatp1a1* expression means that MR images can be obtained longitudinally to track cell migration and growth, and signal intensity can be directly correlated with cell viability. Finally, we and others have found that *Oatp1a1* also enhances the uptake of the firefly luciferase substrate D-luciferin for BLI^43,57^ and the fluorescent dye indocyanine green (using the human ortholog *OATP1B3*) for both fluorescent^58^ and photoacoustic imaging^59^, which gives an added advantage of using *Oatp1* for multi-modality imaging. Since we, and others, have now shown that the human *OATP1B3* gene also functions as a useful fluorescent, photoacoustic and MRI reporter gene, *in vivo*^58,59^, future studies will focus on exchanging *Oatp1a1* for the more translationally favourable *OATP1B3* ortholog.

The improved safety profile and expression of multimodal reporter genes proposed here could have several uses in cell engineering, or at least help answer several concerns with *in vivo* cell therapies. For example, the U.S. Food and Drug Administration have listed potential safety concerns related to unproven stem cell therapies^60^, including: 1) the ability of cells to move from placement sites and change into inappropriate cell types or multiply; 2) failure of cells to work as expected, and 3) the growth of tumours. In addition, the long-term safety profiles of cells engineered with randomly-integrating viruses still require further investigation and optimization. These are concerns that could be addressed by targeting non-viral DNA vectors, such as MCs, to specific safe-harbour loci, such as AAVS1, and reducing the use of integrating viruses. Incorporating reporter genes for clinical grade imaging will also help improve patient safety by allowing one to track cellular therapies *in vivo* (such as for stem cells and immunotherapies). Clinicians could then determine whether the therapeutic cells are localizing to the correct anatomical feature, such as a solid tumour^18^, or to determine their persistence and viability for short- and long-term treatment strategies. Future work will focus on evaluating our system in stem cells and other clinically relevant cell types. Translation will also need to consider building donor vectors that lack optical reporter genes and utilize other selection methods (e.g., magnetic sorting). It is easily feasible to switch out genes from our trimodality construct for other imaging purposes, such as replacing *FLuc2* with a PET reporter gene for dual PET-MR imaging. Suicide switch genes could also be incorporated to further improve safety by killing the engineered cells in cases where they become oncogenic^61^, for example. Not only are these tools useful for clinical cell-based therapies, they are also extremely useful in pre-clinical studies for investigating cancer progression/aggression, metastatic burden and treatment strategies. Avoiding the use of random-integrating viruses and targeted editing should also help reduce off-target effects of gene-editing that may alter the normal characteristics of the cell type being studied.

## Conclusion

Our work demonstrates the first CRISPR/Cas9 HITI MC system for safe harbor integration of a large donor construct encoding three reporter genes for multi-modal longitudinal imaging of cells *in vivo*. We have shown that inclusion of the translationally relevant MR reporter gene, *Oatp1a1*, can enable localization and tracking of small primary and metastatic tumours that are not readily detectable visually or in pre-contrast MR images. This work lays the foundation for an effective and safer genome editing tool for non-invasive reporter gene tracking of multiple cell types *in vivo*, such as for cell-based cancer immunotherapies and stem cell treatments.

## Author Contributions

J.A.R. designed the project, J.A.R, J.J.K. and M.S.M. designed the experiments. J.J.K. directed the study and with M.S.M. carried out most of the experiments. N.N.N. developed the methods for Oatp1a1 MRI. Y.C. helped perform MRI. M.M.E. analyzed MRI data. A.J.H. helped develop the parental plasmids. J.J.K. wrote the manuscript with help from M.S.M., which J.A.R. reviewed and edited.

The authors thank Dr. Kevin Brindle (University of Cambridge) for providing the Oatp1a1 plasmid, Dr. Kristin Chadwick for FACS and flow cytometry help and David Reese for MRI troubleshooting.

## References

1. Brader, P., Serganova, I. & Blasberg, R. G. Noninvasive molecular imaging using reporter genes. Journal of Nuclear Medicine 54, 167–172 (2013).

2. Kircher, M. F., Gambhir, S. S. & Grimm, J. Noninvasive cell-tracking methods. Nat. Rev. Clin. Oncol. 8, 677–688 (2011).

3. Prescher, J. A. & Contag, C. H. Guided by the light: visualizing biomolecular processes in living animals with bioluminescence. Current Opinion in Chemical Biology 14, 80–89 (2010).

4. Hong, H., Yang, Y. & Cai, W. Imaging gene expression in live cells and tissues. Cold Spring Harb. Protoc. 6, (2011).

5. Kim, J. E., Kalimuthu, S. & Ahn, B. C. In Vivo Cell Tracking with Bioluminescence Imaging. Nucl. Med. Mol. Imaging (2010). 49, 3–10 (2015).

6. Li, M., Wang, Y., Liu, M. & Lan, X. Multimodality reporter gene imaging: Construction strategies and application. Theranostics 8, 2954–2973 (2018).

7. Gilad, A. A. & Shapiro, M. G. Molecular Imaging in Synthetic Biology, and Synthetic Biology in Molecular Imaging. Mol. Imaging Biol. 19, 373–378 (2017).

8. Joo, H. K. & Chung, J. K. Molecular-genetic imaging based on reporter gene expression. J. Nucl. Med. 49, 164–180 (2008).

9. Reagan, M. R. & Kaplan, D. L. Concise Review: Mesenchymal Stem Cell Tumor-Homing: Detection Methods in Disease Model Systems. Stem Cells 29, 920–927 (2011).

10. Wang, H. et al. Trafficking Mesenchymal Stem Cell Engraftment and Differentiation in Tumor-Bearing Mice by Bioluminescence Imaging. Stem Cells 27, 1548–1558 (2009).

11. Kidd, S. et al. Direct evidence of mesenchymal stem cell tropism for tumor and wounding microenvironments using in vivo bioluminescent imaging. Stem Cells 27, 2614–2623 (2009).

12. Hamilton, A. M., Parkins, K. M., Murrell, D. H., Ronald, J. A. & Foster, P. J. Investigating the Impact of a Primary Tumor on Metastasis and Dormancy Using MRI: New Insights into the Mechanism of Concomitant Tumor Resistance. Tomography 2, 79–84 (2016).

13. Parkins, K. M. et al. Multimodality cellular and molecular imaging of concomitant tumour enhancement in a syngeneic mouse model of breast cancer metastasis. Sci. Rep. 8, (2018).

14. Parkins, K. M. et al. A multimodality imaging model to track viable breast cancer cells from single arrest to metastasis in the mouse brain. Sci. Rep. 6, (2016).

15. Vandergaast, R. et al. Enhanced noninvasive imaging of oncology models using the NIS reporter gene and bioluminescence imaging. Cancer Gene Therapy (2019). doi: 10.1038/s41417-019-0081-2

16. Shah, K., Jacobs, A., Breakefield, X. O. & Weissleder, R. Molecular imaging of gene therapy for cancer. Gene Therapy 11, 1175–1187 (2004).

17. Yaghoubi, S. S. et al. Noninvasive detection of therapeutic cytolytic T cells with 18 F-FHBG PET in a patient with glioma. Nat. Clin. Pract. Oncol. 6, 53–58 (2009).

18. Keu, K. V. et al. Reporter gene imaging of targeted T cell immunotherapy in recurrent glioma. Sci. Transl. Med. 9, (2017).

19. Milone, M. C. & O’Doherty, U. Clinical use of lentiviral vectors. Leukemia 32, 1529–1541 (2018).

20. Hacein-Bey-Abina, S. et al. Insertional oncogenesis in 4 patients after retrovirus-mediated gene therapy of SCID-X1. J. Clin. Invest. 118, 3132–3142 (2008).

21. Howe, S. J. et al. Insertional mutagenesis combined with acquired somatic mutations causes leukemogenesis following gene therapy of SCID-X1 patients. J. Clin. Invest. 118, 3143–3150 (2008).

22. Hacein-Bey-Abina, S. et al. LMO2-Associated Clonal T Cell Proliferation in Two Patients after Gene Therapy for SCID-X1. Science (80-.). 302, 415–419 (2003).

23. Papapetrou, E. P. & Schambach, A. Gene Insertion Into Genomic Safe Harbors for Human Gene Therapy. Mol. Ther. 24, 678–684 (2016).

24. Wang, Y. et al. Genome editing of human embryonic stem cells and induced pluripotent stem cells with zinc finger nucleases for cellular imaging. Circ. Res. 111, 1494–1503 (2012).

25. Luo, Y. et al. Stable Enhanced Green Fluorescent Protein Expression After Differentiation and Transplantation of Reporter Human Induced Pluripotent Stem Cells Generated by AAVS1 Transcription Activator-Like Effector Nucleases. Stem Cells Transl. Med. 3, 821–835 (2014).

26. Cerbini, T. et al. Transcription activator-like effector nuclease (TALEN)-mediated CLYBL targeting enables enhanced transgene expression and one-step generation of dual reporter human induced pluripotent stem cell (iPSC) and neural stem cell (NSC) lines. PLoS One 10, (2015).

27. Cho, S. W., Kim, S., Kim, J. M. & Kim, J. S. Targeted genome engineering in human cells with the Cas9 RNA-guided endonuclease. Nat. Biotechnol. 31, 230–232 (2013).

28. Cong, L. et al. Multiplex genome engineering using CRISPR/Cas systems. Science (80-.). 339, 819–823 (2013).

29. Jinek, M. et al. RNA-programmed genome editing in human cells. Elife 2013, (2013).

30. Mali, P. et al. RNA-guided human genome engineering via Cas9. Science (80-.). 339, 823–826 (2013).

31. Ding, Q. et al. Enhanced efficiency of human pluripotent stem cell genome editing through replacing TALENs with CRISPRs. Cell Stem Cell 12, 393–394 (2013).

32. Dubois, V. P. et al. Safe Harbor Targeted CRISPR-Cas9 Tools for Molecular-Genetic Imaging of Cells in Living Subjects. Cris. J. 1, 440–449 (2018).

33. Orthwein, A. et al. A mechanism for the suppression of homologous recombination in G1 cells. Nature 528, 422–426 (2015).

34. Lieber, M. R. The Mechanism of Double-Strand DNA Break Repair by the Nonhomologous DNA End-Joining Pathway. Annu. Rev. Biochem. 79, 181–211 (2010).

35. Auer, T. O., Duroure, K., De Cian, A., Concordet, J. P. & Del Bene, F. Highly efficient CRISPR/Cas9-mediated knock-in in zebrafish by homology-independent DNA repair. Genome Res. 24, 142–153 (2014).

36. Suzuki, K. & Izpisua Belmonte, J. C. In vivo genome editing via the HITI method as a tool for gene therapy. J. Hum. Genet. 63, 157–164 (2018).

37. He, X. et al. Knock-in of large reporter genes in human cells via CRISPR/Cas9-induced homology-dependent and independent DNA repair. Nucleic Acids Res. 44, (2016).

38. Suzuki, K. et al. In vivo genome editing via CRISPR/Cas9 mediated homology-independent targeted integration. Nature 540, 144–149 (2016).

39. Wang, T. D., Chen, Y. & Ronald, J. A. A novel approach for assessment of prostate cancer aggressiveness using survivin-driven tumour-activatable minicircles. Gene Ther. 26, 177–186 (2019).

40. Ronald, J. A. et al. Development and Validation of Non-Integrative, Self-Limited, and Replicating Minicircles for Safe Reporter Gene Imaging of Cell-Based Therapies. PLoS One 8, (2013).

41. Darquet, A. M., Cameron, B., Wils, P., Scherman, D. & Crouzet, J. A new DNA vehicle for nonviral gene delivery: Supercoiled minicircle. Gene Ther. 4, 1341–1349 (1997).

42. Patrick, P. S. et al. Dual-modality gene reporter for in vivo imaging. Proc. Natl. Acad. Sci. U. S. A. 111, 415–20 (2014).

43. Nyström, N. N. et al. Longitudinal Visualization of Viable Cancer Cell Intratumoral Distribution in Mouse Models Using Oatp1a1-Enhanced Magnetic Resonance Imaging. Invest. Radiol. 54, 302–311 (2019).

44. Yushkevich, P. A. et al. User-guided 3D active contour segmentation of anatomical structures: Significantly improved efficiency and reliability. Neuroimage 31, 1116–1128 (2006).

45. Paix, A. et al. Precision genome editing using synthesis-dependent repair of Cas9-induced DNA breaks. Proc. Natl. Acad. Sci. U. S. A. 114, E10745–E10754 (2017).

46. Kreiss, P. et al. Plasmid DNA size does not affect the physicochemical properties of lipoplexes but modulates gene transfer efficiency. Nucleic Acids Res. 27, 3792–3798 (1999).

47. Hornstein, B. D., Roman, D., Arévalo-Soliz, L. M., Engevik, M. A. & Zechiedrich, L. Effects of circular DNA length on transfection efficiency by electroporation into HeLa cells. PLoS One 11, (2016).

48. Pellenz, S. et al. New Human Chromosomal Sites with ‘Safe Harbor’ Potential for Targeted Transgene Insertion. Hum. Gene Ther. 30, 814–828 (2019).

49. Ordovás, L. et al. Efficient recombinase-mediated cassette exchange in hPSCs to study the hepatocyte lineage reveals AAVS1 locus-mediated transgene inhibition. Stem Cell Reports 5, 918–931 (2015).

50. Zhang, X. H., Tee, L. Y., Wang, X. G., Huang, Q. S. & Yang, S. H. Off-target effects in CRISPR/Cas9-mediated genome engineering. Molecular Therapy - Nucleic Acids 4, e264 (2015).

51. Wang, D. C. & Wang, X. Off-target genome editing: A new discipline of gene science and a new class of medicine. Cell Biology and Toxicology 35, 179–183 (2019).

52. Li, P. et al. Cas12a mediates efficient and precise endogenous gene tagging via MITI: microhomology-dependent targeted integrations. Cell. Mol. Life Sci. (2019). doi: 10.1007/s00018-019-03396-8

53. Vakulskas, C. A. et al. A high-fidelity Cas9 mutant delivered as a ribonucleoprotein complex enables efficient gene editing in human hematopoietic stem and progenitor cells. Nat. Med. 24, 1216–1224 (2018).

54. Chen, J. S. et al. Enhanced proofreading governs CRISPR-Cas9 targeting accuracy. Nature 550, 407–410 (2017).

55. Oliveira, P. H. & Mairhofer, J. Marker-free plasmids for biotechnological applications - implications and perspectives. Trends in Biotechnology 31, 539–547 (2013).

56. Xiao, Y. D. et al. MRI contrast agents: Classification and application (Review). International Journal of Molecular Medicine 38, 1319–1326 (2016).

57. Patrick, P. S., Lyons, S. K., Rodrigues, T. B. & Brindle, K. M. Oatp1 Enhances Bioluminescence by Acting as a Plasma Membrane Transporter for d-luciferin. Mol. Imaging Biol. 16, 626–634 (2014).

58. Wu, M. R. et al. Organic anion-transporting polypeptide 1B3 as a dual reporter gene for fluorescence and magnetic resonance imaging. FASEB J. 32, 1705–1715 (2018).

59. Nyström, N. N., Yip, L. C. M., Carson, J. J. L., Scholl, T. J. & Ronald, J. A. Development of a Human Photoacoustic Imaging Reporter Gene Using the Clinical Dye Indocyanine Green. Radiol. Imaging Cancer 1, e190035 (2019).

60. Marks, P. W., Witten, C. M. & Califf, R. M. Clarifying Stem-Cell Therapy’s Benefits and Risks. N. Engl. J. Med. 376, 1007–1009 (2017).

61. Ivics, Z. Self-Destruct Genetic Switch to Safeguard iPS Cells. Molecular Therapy 23, 1417–1420 (2015).

