## Supplemental Figures for "A Safe Harbor-Targeted CRISPR/Cas9 Homology Independent Targeted Integration (HITI) System for Multi-Modality Reporter Gene-Based Cell Tracking"

#### Supplemental Figure 1

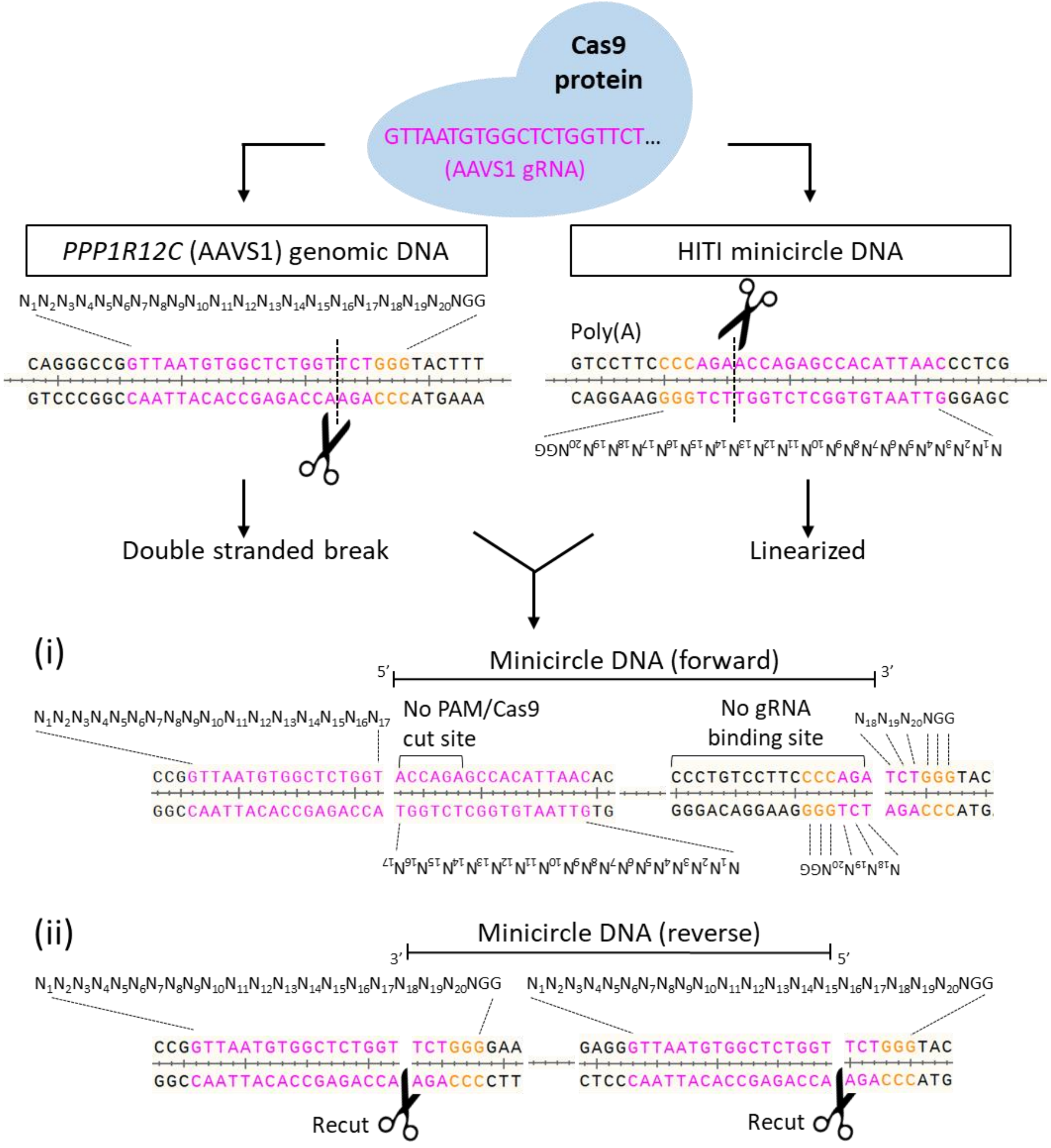


Supplemental Figure 1. HITI sequence at the AAVS1 locus and reciprocal sequence in our HITI minicircle. Cas9/gRNA induce double stranded DNA breaks and linearization of the minicircle. The two possible integration orientations are shown. The Cas9 cut sites are lost in the forward direction (i), whereas they are preserved in the reverse integration (ii) and should then be recut until the donor minicircle DNA is inserted in the correct (forward) direction.

### Supplemental Figure 2

**A** 1st tdT+ / zsG+ sort

| | % tdT+ | % zsG+ | % tdT+, zsG+ | % Purity |
|---|---|---|---|---|
| HDR AAVS1 | 71.7 | 67.8 | 40.1 | 88.9 |
| HDR SCR | 69.1 | 71.2 | 41 | 89.2 |
| HITI AAVS1 | 73.3 | 77.5 | 47.5 | 95.2 |
| HITI SCR | 69.5 | 74 | 39.5 | 93.6 |

| | % tdT+ | % zsG+ | % tdT+, zsG+ | % Purity |
|---|---|---|---|---|
| HDR AAVS1 | 47.6 | 47.1 | 23.8 | 93.7 |
| HDR SCR | 35.7 | 35.1 | 17.3 | 95.2 |
| HITI AAVS1 | 39.5 | 46 | 23.2 | 94.9 |
| HITI SCR | 40.5 | 52.4 | 22.5 | 90.2 |

| | % tdT+ | % zsG+ | % tdT+, zsG+ | % Purity |
|---|---|---|---|---|
| HDR AAVS1 | 28.9 | 36.9 | 26.3 | 96.7 |
| HDR SCR | 36.5 | 50.8 | 33.9 | 96.4 |
| HITI AAVS1 | 38.3 | 43.2 | 35.1 | 95.1 |
| HITI SCR | 37.7 | 46.9 | 35 | 97.3 |

| | % tdT+ | % zsG+ | % tdT+, zsG+ | % Purity |
|---|---|---|---|---|
| HDR AAVS1 | 14.6 | 18 | 13.5 | 96.3 |
| HDR SCR | 11.2 | 14.2 | 10.4 | 97.5 |
| HITI AAVS1 | 11.3 | 15.7 | 10.4 | 90.4 |
| HITI SCR | 11.1 | 16.4 | 10.2 | 92.1 |

| | % tdT+ | % zsG+ | % tdT+, zsG+ | % Purity |
|---|---|---|---|---|
| HDR AAVS1 | 9.3 | 15.5 | 10.3 | 89.6 |
| HDR SCR | 5.0 | 4.2 | 4.9 | 92.5 |
| HITI AAVS1 | 6.3 | 7.9 | 6.6 | 89.3 |
| HITI SCR | 5.9 | 12.3 | 6.7 | 94.8 |

| | % tdT+ | % zsG+ | % tdT+, zsG+ | % Purity |
|---|---|---|---|---|
| HDR AAVS1 | 3.6 | 4.2 | 3.9 | 91.8 |
| HDR SCR | 7.7 | 13.8 | 8.6 | 97.8 |
| HITI AAVS1 | 3.1 | 3.4 | 3.3 | 92.6 |
| HITI SCR | 4.5 | 9.7 | 5.3 | 92.3 |

**B**

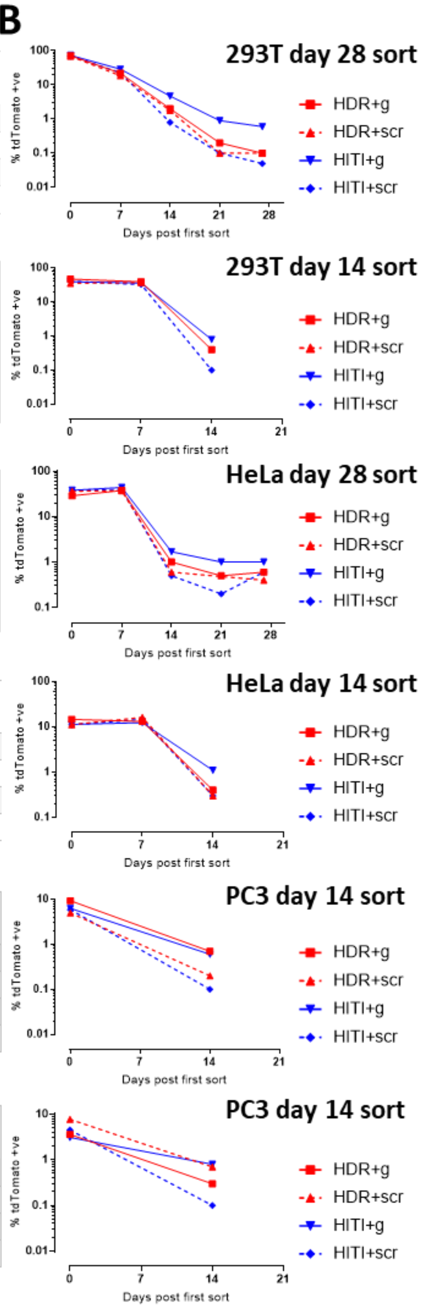


**C** 2nd tdT+ / zsG- sort

| | % tdT+ | % tdT+, zsG- |
|---|---|---|
| HDR AAVS1 | 0.1 | 0.1 |
| HDR SCR | 0.1 | 0.1 |
| HITI AAVS1 | 0.6 | 0.6 |
| HITI SCR | 0 | 0 |

| | % tdT+ | % tdT+, zsG- |
|---|---|---|
| HDR AAVS1 | 0.4 | 0.3 |
| HDR SCR | n/a | n/a |
| HITI AAVS1 | 0.8 | 0.8 |
| HITI SCR | 0.1 | 0.1 |

| | % tdT+ | % tdT+, zsG- |
|---|---|---|
| HDR AAVS1 | 0.7 | 0.6 |
| HDR SCR | 0.6 | 0.4 |
| HITI AAVS1 | 1.1 | 1 |
| HITI SCR | 0.9 | 0.6 |

| | % tdT+ | % tdT+, zsG- |
|---|---|---|
| HDR AAVS1 | 0.4 | 0.3 |
| HDR SCR | 0.3 | 0.1 |
| HITI AAVS1 | 1.1 | 0.9 |
| HITI SCR | 0.3 | 0.1 |

| | % tdT+ | % tdT+, zsG- |
|---|---|---|
| HDR AAVS1 | 1.5 | 0.7 |
| HDR SCR | 0.7 | 0.2 |
| HITI AAVS1 | 1.1 | 0.6 |
| HITI SCR | 0.6 | 0.1 |

| | % tdT+ | % tdT+, zsG- |
|---|---|---|
| HDR AAVS1 | 1.1 | 0.3 |
| HDR SCR | 1.4 | 0.7 |
| HITI AAVS1 | 1.8 | 0.8 |
| HITI SCR | 1.6 | 0.1 |

Supplemental Figure 2. FACS of 293T, HeLa and PC3 cells. *A*, The percentage of tdTomato (tdT+) and zsGreen (zsG+) cells that were sorted 48 hours after transfection is shown (double positives shaded in brown). *B*, tdT fluorescence was tracked every 7 days with flow cytometry. *C*, Cells were sorted for tdT+ fluorescence *only* for each cell line at 14- or 21-days post 1st sort (shaded in pink). n/a, 293T HDR-SCR cells were lost due to infection prior to final sort.

#### Supplemental Figure 3

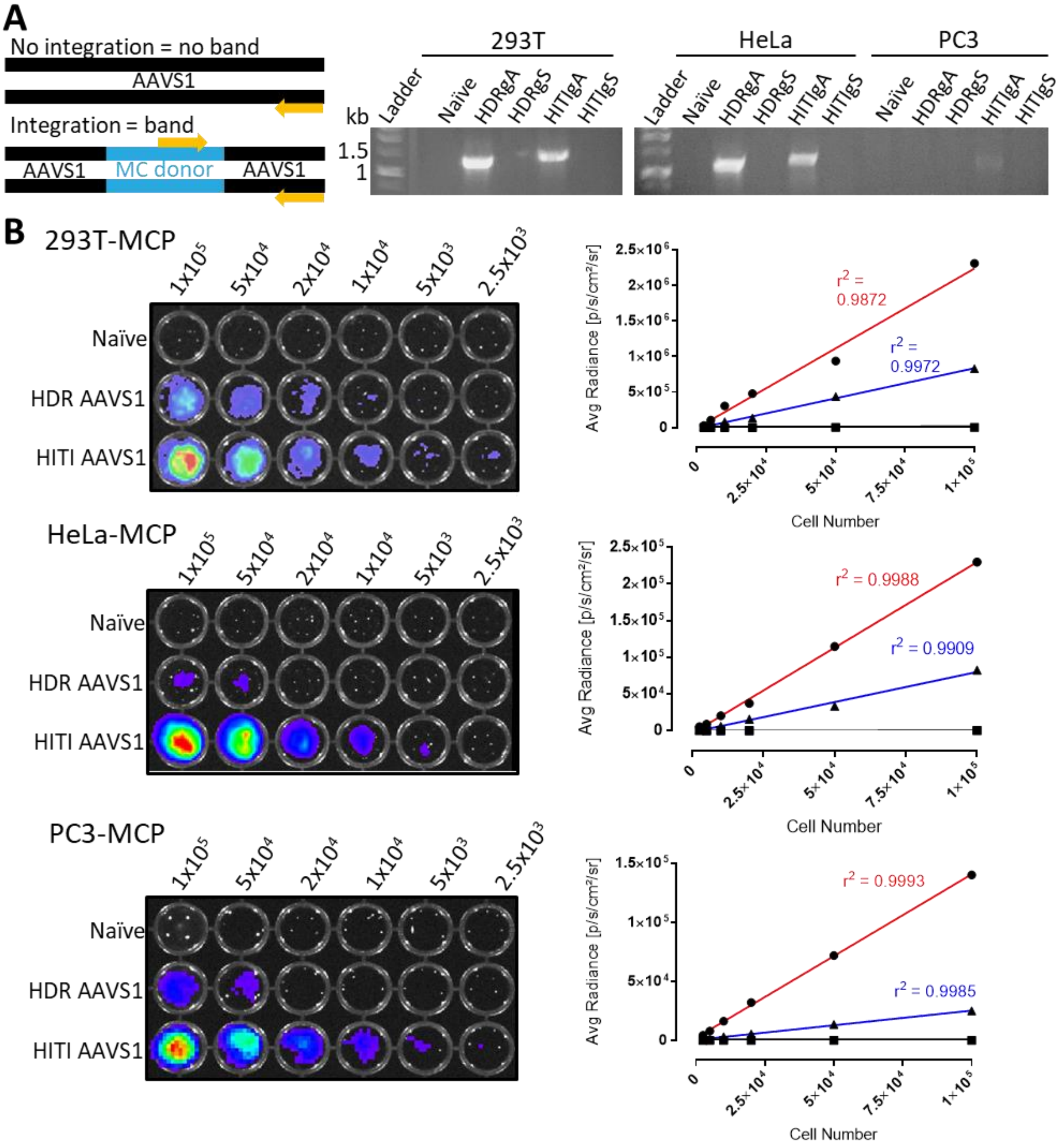


Supplemental Figure 3. Integration and *FLuc2* reporter gene checks for MCPs. *A*, Forward and reverse primers were designed to amplify junctional bands of 1.4 kb for HITI and 1.3 kb for HDR engineered cells with correct MC integration. PCR gel electrophoresis showed positive integration bands for all cell types engineered with HITI-AAVS1-gRNA and for 293T and HeLa cells engineered with HDR-AAVS1-gRNA. *B*, BLI analysis of MCPs with increasing cell numbers for HDR and HITI engineered cells.

### Supplemental Figure 4

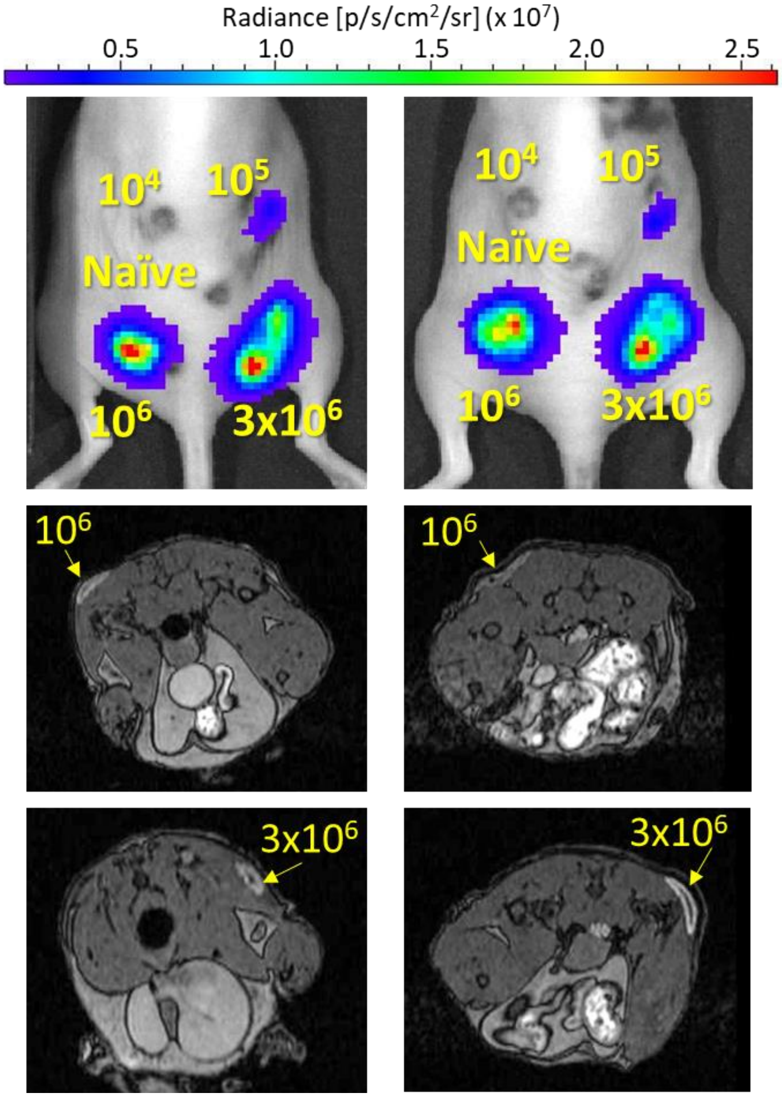


Supplemental Figure 4. BLI and Oatp1a1 sensitivity *in vivo*. Remaining two mice from n = 3 cohort shown side-by-side with indicated numbers of PC3-HITI cells injected in yellow (same experiment as shown in Figure 4). $1^{st}$ row, BLI; 2nd-$3^{rd}$ rows, representative MRI of enhanced contrast in regions of $10^6$ and $3x10^6$ PC3-HITI cells.

### Supplemental Figure 5

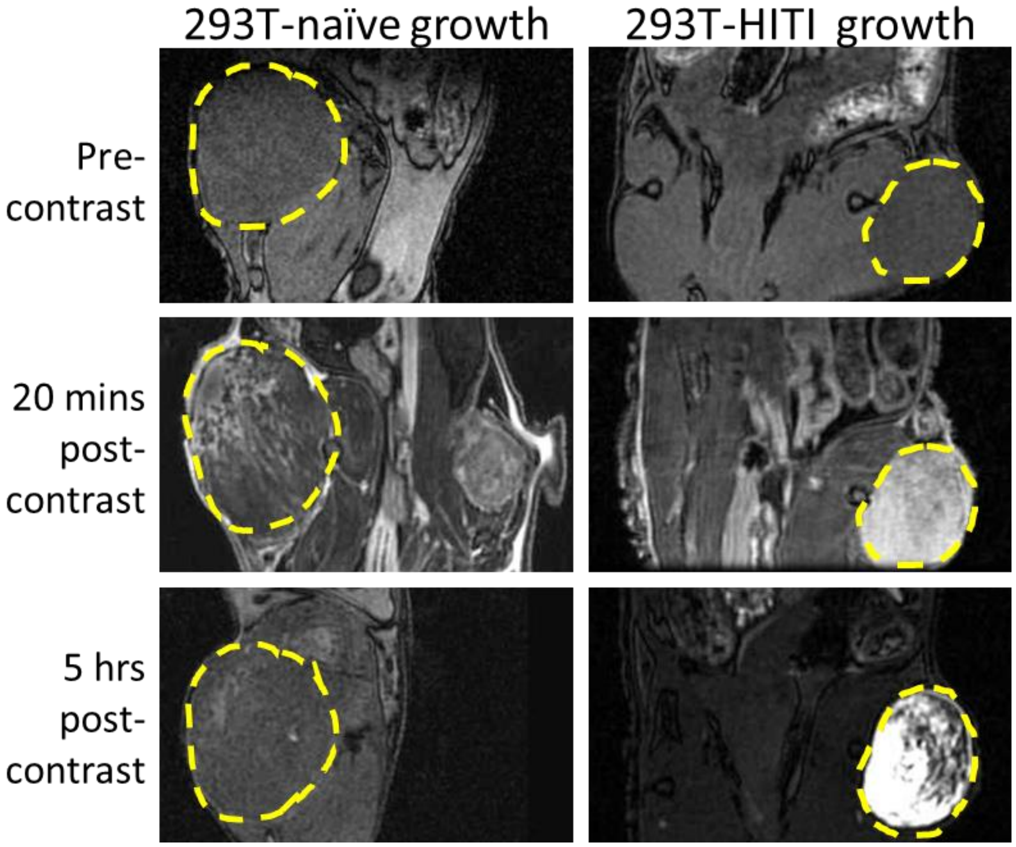


Supplemental Figure 5. *In vivo* MRI of subcutaneous 293-HITI clone growths. Palpable tumours were evident for PC3 naïve and HITI clonal cells on the left and right flanks of mice in pre-contrast images (delineated with dashed yellow lines). Increased contrast was evident in both naïve and HITI cells 20 mins post Gd-EOB-DTPA injection. At 5 hrs post contrast injection, only the HITI engineered cells retained the Gd-EOB-DTPA agent.

Supplemental Figure 6

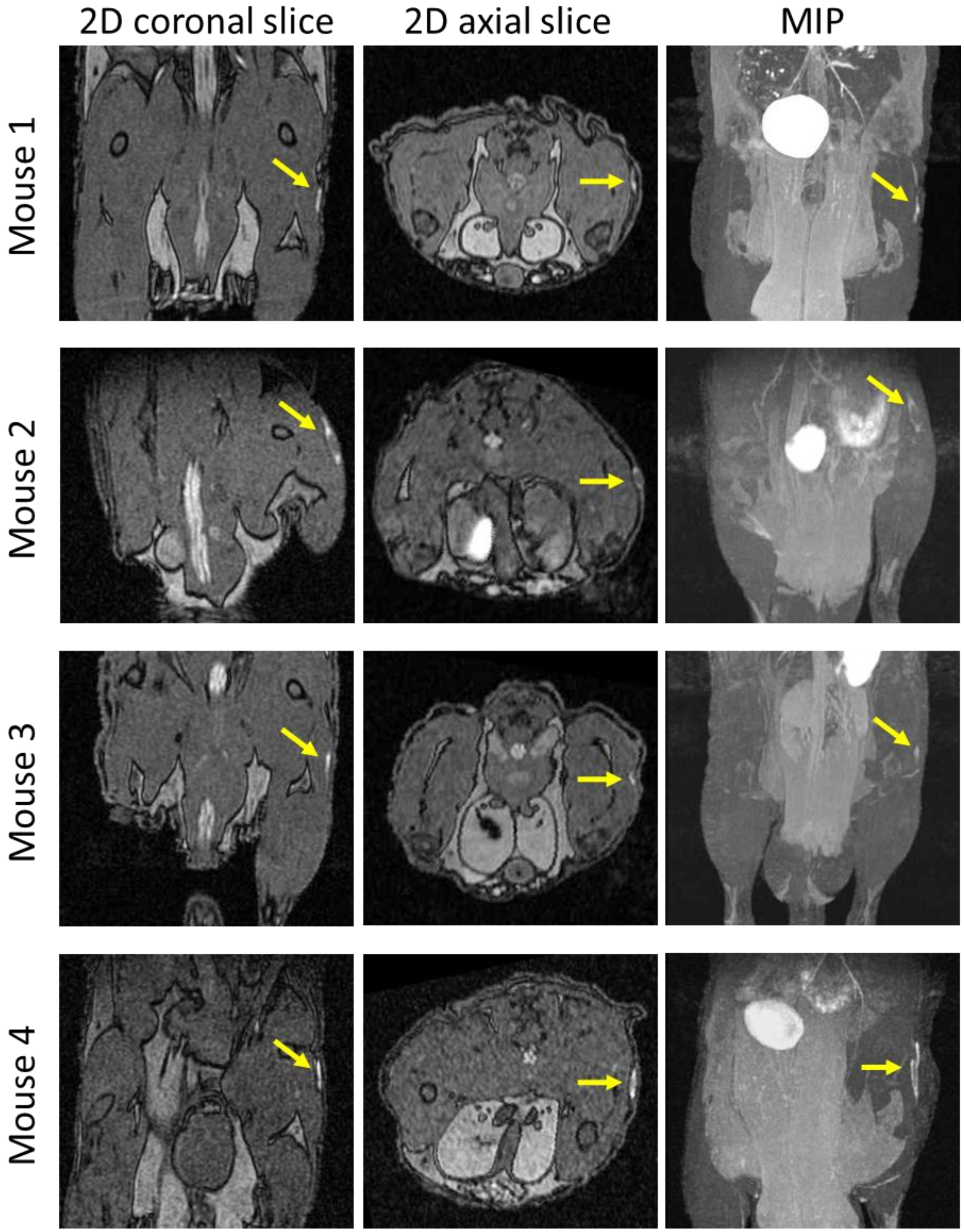


Supplemental Figure 6. Subcutaneous Oatp1a1 expressing tumours at day 11 post injection. Images shown are from MRI scans performed 5 hours after Gd-EOB-DTPA injection. Yellow arrows indicate positive contrast from Oatp1a1-expressing PC3-HITI subcutaneous tumours. MIP, maximum intensity projection.

Supplemental Figure 7

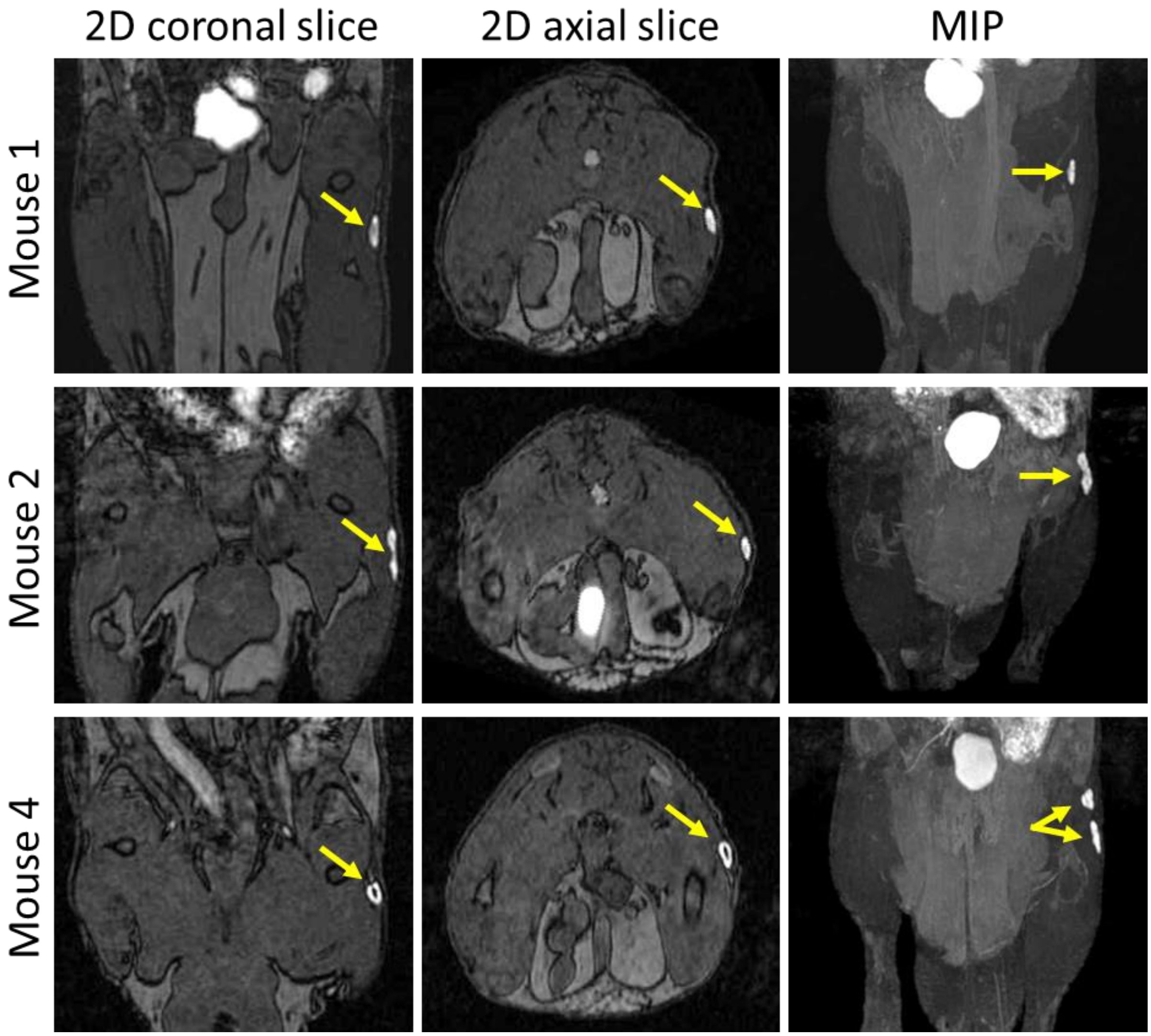


Supplemental Figure 7. Subcutaneous Oatp1a1 expressing tumours at day 46 post injection. Images shown are from MRI scans performed 5 hours after Gd-EOB-DTPA injection. Yellow arrows indicate positive contrast from Oatp1a1-expressing PC3-HITI subcutaneous tumours. N.B. Mouse 3 was sacked part way through the study due to an unrelated health condition. MIP, maximum intensity projection.
